# Transcriptome Dynamics Reveals Progressive Transition from Effector to Memory in CD4^+^ T cells

**DOI:** 10.1101/675967

**Authors:** Megan S. F. Soon, Hyun Jae Lee, Jessica A. Engel, Jasmin Straube, Bryce S. Thomas, Lachlan S. Clarke, Pawat Laohamonthonkul, Clara P. S. Pernold, Rohit N. Haldar, Cameron G. Williams, Lianne I. M. Lansink, Ross Koufariotis, Vanessa Lakis, Scott Wood, Xi Chen, Kylie R. James, Tapio Lönnberg, Steven W. Lane, Miles P. Davenport, David S. Khoury, Valentine Svensson, Sarah A. Teichmann, Ashraful Haque

## Abstract

CD4^+^ T cells are repositories of immune memory, conferring enhanced immunity to many infectious agents. Studies of acute viral and bacterial infection suggest that memory CD4^+^ T cells develop directly from effectors. However, delineating these dynamic developmental pathways has been challenging. Here, we used high-resolution single-cell RNA-seq and temporal mixture modelling to examine the fate of Th1 and Tfh effector cells during non-lethal *Plasmodium* infection in mice. We observed linear Th1 and Tfh pathways towards memory, characterized by progressive halving in the numbers of genes expressed, and partial transcriptomic coalescence. Low-level persisting infection diverted but did not block these pathways. We observed in the Th1-pathway a linear transition from Th1 through a Tr1 state to T_EM_ cells, which were then poised for Th1 re-call. The Tfh-pathway exhibited a modest Th1-signature throughout, with little evidence of Tr1 development, and co-expression of T_CM_ and memory Tfh markers. Thus, we present a high-resolution atlas of transcriptome dynamics for naïve to memory transitions in CD4^+^ T cells. We also defined a subset of memory-associated genes, including transcription factors *Id2* and *Maf*, whose expression increased progressively against the background of transcriptomic quiescence. Single-cell ATAC-seq revealed substantial heterogeneity in chromatin accessibility in single effectors, which was extensively, though incompletely reset and homogenized in memory. Our data reveal that linear transitions from effector to memory occur in a progressive manner over several weeks, suggesting opportunities for manipulating CD4^+^ T cell memory after primary infection.

**Highlights:**

- scRNA-seq reveals progressive transition from effector to memory in CD4^+^ T cells.
- Transcriptome dynamics suggest linear not branching models for memory development.
- A subset of genes associates with gradual onset of CD4^+^ T cell memory.
- Th1/Tfh predisposition varies among clonotypes with identical antigen-specificity.
- scATAC-seq uncovers non-coding “memory” elements in the genome.

## Introduction

CD4^+^ T cells are important repositories of immunological memory, and confer enhanced immunity to many infectious agents. They can be classified into circulating effector memory (T_EM_) and central memory (T_CM_) cells (Sallusto et al., 1999), or non-circulating tissue-resident memory (T_RM_) cells (Hogan et al., 2001; Masopust and Soerens, 2019). Memory CD4^+^ T cells are not produced *de novo* in primary lymphoid organs, but result from peptide:MHCII-mediated priming of individual naïve CD4^+^ T cells in secondary lymphoid tissue. Primed CD4^+^ T cells undergo clonal expansion, and differentiate into a wide spectrum of effector T helper (Th) phenotypes, including Th1, Th2, Th17, T follicular helper (Tfh) and iTreg cells (Taniuchi, 2018). After a period of contraction, a memory population is detected, that is quantitatively more prevalent than naïve precursors, and is qualitatively enhanced in recall capacity. One hypothesis from this prevailing model is that memory CD4^+^ T cells develop directly from effector precursors. A number of approaches have been employed to test this, including fate mapping of cytokine or chemokine receptor gene expression (Harrington et al., 2008; Pepper et al., 2011), adoptive transfer of cell-sorted effector sub-populations (Hale et al., 2013; Marshall et al., 2011), and single-cell transfer and longitudinal sampling (Tubo et al., 2016). Data from these studies suggest that if a single naïve CD4^+^ T cell produces effector progeny, it will also give rise to a memory cell population that partly resembles the previous effector population. This scenario is consistent with both a linear model of differentiation (i.e. naïve to effector to memory states), as well as a branching model, in which memory lineages develop before or during effector differentiation. Since these models are not mutually exclusive, the extent to which they might co-exist in different infection scenarios is unclear. Marking effector subsets to assess their potential transition to memory has mainly focused on a small number of molecules including IFNγ, CCR7, CXCR5, PSGL1 and Ly6C (Hale et al., 2013; Harrington et al., 2008; Marshall et al., 2011; Pepper et al., 2011; Tubo et al., 2016). Using these markers in acute systemic bacterial and viral infection models, Th1 effectors appeared to give rise to Th1-poised T_EM_ cells. Similarly, cells referred to as either Tfh or T_CM_ precursor cells, appeared to give rise to T_CM_ cells. A recent study employing single-cell transcriptomics during LCMV-infection reported transcriptional differences existed between Tfh and T_CM_ precursor cells (Ciucci et al., 2019). Here, we employed an established protozoan parasitic infection model in combination with single-cell transcriptomics and computational modelling to monitor CD4^+^ T cells over extended periods of time (James et al., 2018; Lonnberg et al., 2017). We captured transcriptomes of antigen-specific CD4^+^ T cells from *Plasmodium-*infected mice that spanned naïve, effector, regulatory, exhausted and memory states. Then, by performing temporal mixture modeling of the single-cell RNA-sequencing (scRNA-seq) data, and comparison to single-cell epigenomic data, we explored whether transcriptional dynamics and chromatin accessibility could capture developmental links between effector and memory states.

## Results

### PbTII cells exhibit heterogeneous effector fates and persist for extended periods of time

To explore the fate of effector CD4^+^ T cell responses *in vivo*, we first assessed a protozoan parasitic infection model, *Plasmodium chabaudi chabaudi* AS (*Pc*AS) infection of C57BL/6J mice (Suss et al., 1988). This infection model is characterized by peak parasite load by ∼1 week, as well as splenomegaly and a low-level persisting parasite burden for ∼2-3 months. Mice exhibit a protective CD4^+^ T cell response dominated by Th1 and Tfh cells (Meding et al., 1990; Perez-Mazliah et al., 2015). Given that TCR sequence influences effector fate decisions (Tubo et al., 2013), we controlled for this using PbTII cells (Fernandez-Ruiz et al., 2017), a CD4^+^ TCR transgenic T cell line that differentiates simultaneously into T-bet^hi^ IFNγ^+^ Th1 cells and Bcl6^hi^ CXCR5^hi^ Tfh effector cells by the end of the first week of *Pc*AS infection (Lonnberg et al., 2017), but whose long term kinetics were unknown. The PbTII/*Pc*AS model permitted tracking of two independent CD4^+^ T cell effector types in one infection. In addition, to mitigate the potential for persisting parasitic infection to trigger T cell exhaustion and impair memory onset, we treated mice from peak with the anti-malarial drug, sodium artesunate, or control saline (Figure S1).

After transfer of a physiological number of naïve PbTIIs and infection with *Pc*AS, splenic PbTII numbers peaked at day 7, and remained detectable well beyond the first month of infection (Figure 1A). This remained the case even after treatment from peak with the anti-malarial, sodium artesunate (Figure S1A-C), although the drug induced further reduction in PbTII numbers over time. Splenic PbTIIs displayed heterogeneous CD62L expression during the first month of infection, which was partly reduced under drug treatment (Figure 1B). Bifurcated expression of Th1-associated, CXCR6, and Tfh-associated, CXCR5 at peak was retained beyond the first month of infection (Figure 1C), although decreasing proportions of cells expressed these receptors over time. Amongst control-treated (Figure 1D) or artesunate-treated mice (Figure 1E), bifurcated CXCR5/6 expression was apparent on both CD62L^hi^ and CD62L^lo^ populations. Together these data revealed substantial heterogeneity and sub-populations of splenic PbTIIs, even when assessment was confined to just three markers. Heterogeneity was also evident in micro-anatomical location, since effector PbTIIs were variously located in splenic T cell, B cell and red pulp zones at day 7 *p.i.* (Figure S2A-B), as well as in germinal centres and B cell zones at later timepoints (Figure S2C-D). Our data indicated that PbTIIs entered germinal centre reactions, and persisted long enough to potentially become memory CD4^+^ T cells.

**Figure 1.**
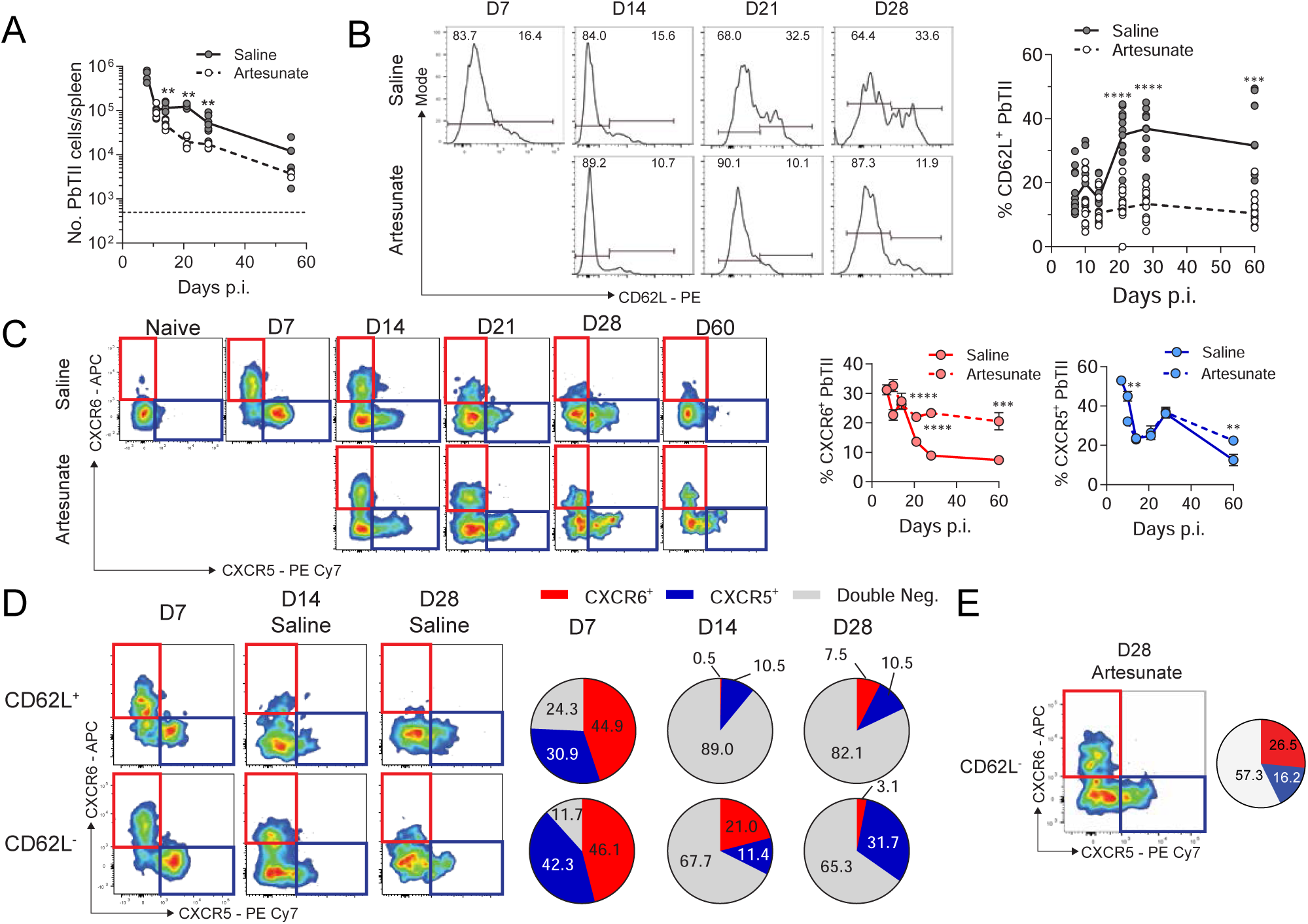
Characterization of memory development in PbTII cells during *in vivo Plasmodium* infection. **(A)** Changes in splenic PbTII cell numbers during infection with *Pc*AS. **(B)** Representative fluorescence-activated cell sorting (FACS) plots for surface CD62L expression on PbTII cells. Graph of proportion of CD62L^+^ PbTII cells over time. **(C)** Representative FACS plots for surface CXCR6 and CXCR5 expression on PbTII cells. Graph of proportion of CXCR6^+^ or CXCR5^+^ PbTII cells over time. **(D)** Representative FACS plots for CXCR5 and CXCR6 expression on either CD62L^+^ or CD62L^−^ PbTII cells from saline control group. Pie charts of proportion of CXCR6^+^, CXCR5^+^ or double negative PbTII cells derived from either CD62L^+^ or CD62L^−^ populations, from indicated timepoints. **(E)** Representative FACS plot and pie chart as in (D) for CD62L^−^ PbTII cells from artesunate-treated mice at D28 *p.i.* Data for CD62L^+^ cells was excluded due to the low number of cells present for analysis. All data is pooled from 2-3 independent experiments (n=5-6 mice/ group/ timepoint). Statistical analysis was performed between saline- and artesunate-treated groups for each individual timepoint using Mann-Whitney test. Each circle represents an individual mouse (A-B), while data is presented as mean +/- SEM (C). *p< 0.05, **p< 0.01, ***p<0.001, ****p<0.0001

### PbTIIs at late timepoints exhibit altered chromatin structure and exhibit recall responses during homologous re-challenge

We next determined whether PbTIIs present after four weeks of infection exhibited memory responses. While direct *ex vivo* production of the Th1 cytokine IFNγ progressively waned from day 7 to day 28 *p.i*. (Figure 2A & S3A), the capacity to mount a recall response, as judged by *in vitro* re-stimulation was evident throughout (Figure 2B & S3B), and was substantially enhanced when persisting infection was removed by artesunate treatment. The potential for PbTIIs to mount recall responses that resembled Th1/Tfh effector responses was also supported by assessment of chromatin accessibility in pooled cells via bulk ATAC-seq (Figure 2C-D). We noted that genome-wide regions of accessible chromatin found at day 7 *p.i.* were not as prevalent in naïve PbTIIs, but were partially evident at day 28 *p.i.* regardless of whether mice were drug treated or not (Figure 2C). This was also seen for specific Th1/Tfh associated genes, *Ifng*, *Cxcr5*, *Tbx21* and *Il21* (Figure 2D). These bulk assessments suggested PbTIIs present at the end of the month had partially retained effector chromatin structure, suggesting some potential to mount recall Th1 and Tfh recall responses *in vivo.* To test this, we performed homologous high-dose re-challenge in mice harbouring *in vivo* primed PbTIIs for 28 days, with naïve, congenically marked naïve PbTIIs co-transferred the day before re-challenge as comparators (Figure 2E). Consistent with *in vitro* data (Figure 2B), PbTIIs primed 28 days previously, exhibited greater direct *ex vivo* IFNγ production compared to co-transferred naïve PbTIIs. Notably, PbTIIs in artesunate-treated mice exhibited a substantially greater Th1 recall response and CD69 upregulation, but lower Ki67 expression, compared to PbTIIs in control saline treated mice (Figure 2F & S3C-D). Together these data confirmed that amongst PbTIIs present 4 weeks after initial priming, particularly if infection was controlled with anti-malarial drugs, were those capable of mounting a potent Th1 recall response *in vivo*, justifying our subsequent classification of them as T_EM_ cells with a Th1-bias.

**Figure 2.**
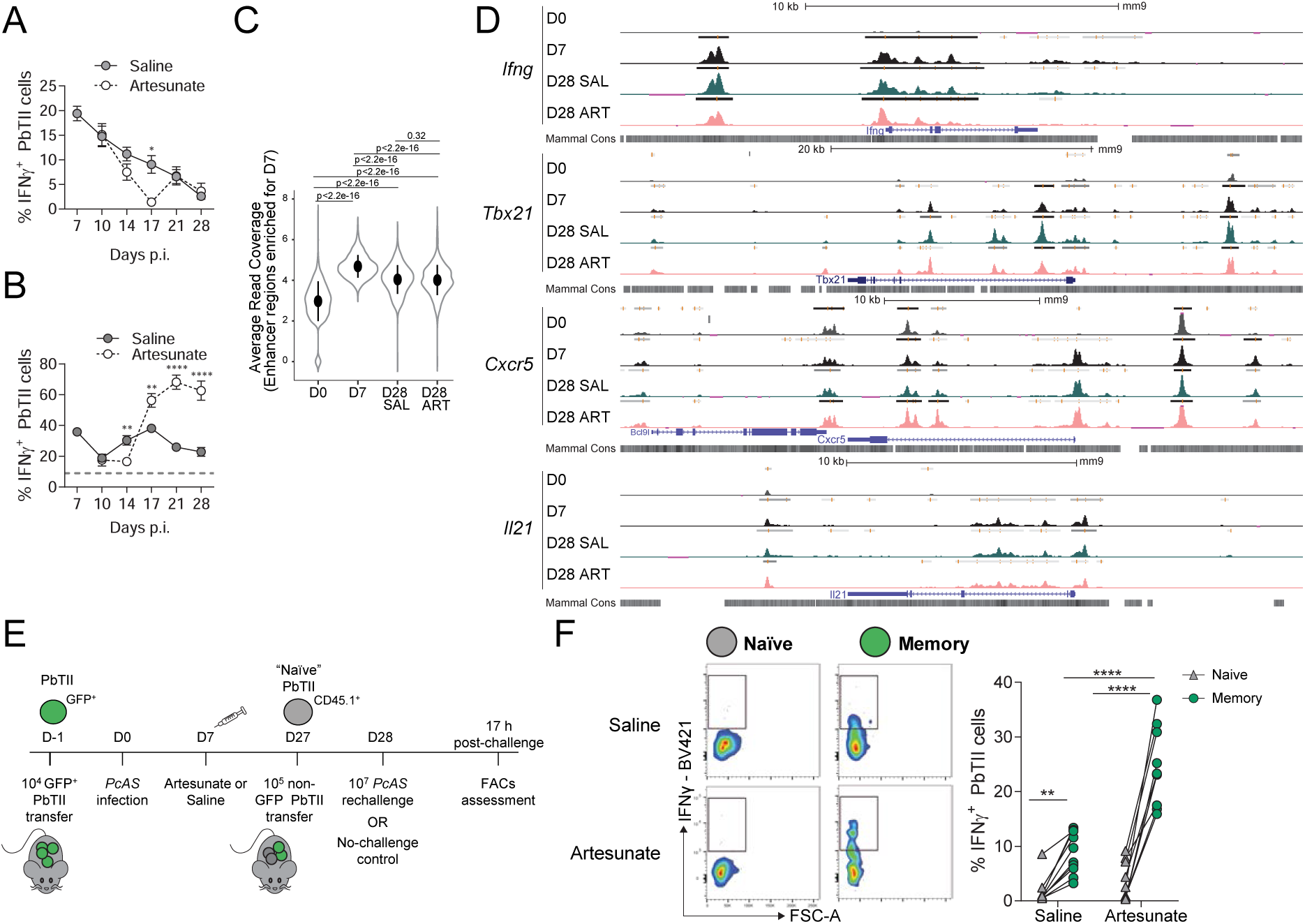
PbTII cells exhibit functional memory characteristics. **(A)** Analysis of *ex vivo* IFNγ production by PbTII cells in the absence of any restimulation. **(B)** Analysis of *ex vivo* IFNγ production by PbTII cells after restimulation with PMA and ionomycin for 3 hours *in vitro*. Dashed grey line represents threshold of IFNγ production by naïve PbTII cells. **(C)** Average number of reads for ATAC-seq peaks within enhancer regions enriched for D7 *p.i.* sample. Bulk ATAC-seq experiments were performed as 2 independent experiments. Data is derived using overlapping peaks shared between biological replicates from each experiment for individual timepoints. **(D)** Mean ATAC-seq peak coverage at *Ifng*, *Tbx21*, *Cxcr5* and *Il21* gene loci with a scale of 0-45 for all tracks. Boxes represent peaks called using MACS2. Data is shown for a representative experiment out of 2 independent biological repeats. **(E)** Schematic for model for studying PbTII responses during *in vivo* rechallenge (refer to Methods for details). **(F)** Representative FACS plots showing *ex vivo* production of IFNγ without any further re-stimulation at 17 hours post-rechallenge for memory (green) or naïve (grey) PbTII cells. Graph comparing IFNγ production between memory or naïve PbTII cells for saline- or artesunate-treated group. Data is pooled from 2 independent experiments (n=5-6 mice / group/ timepoint). Statistical analysis was performed between saline- and artesunate-treated group for each individual timepoints using Mann-Whitney test (A-B), or using paired analysis (F). (A-B, F) *p< 0.05, **p< 0.01, ***p<0.001, ****p<0.0001. Each dot represents an individual mouse (F), while data is presented as mean+/- SEM (A-B). Sample comparison was calculated using Wilcoxon-rank test (C).

### PbTIIs exhibit only two effector fates during infection, whose transcriptomes change over time and in response to anti-malarial drugs

Since PbTIIs were retained for extended periods of time and exhibited recall responses, we next sought to monitor the molecular phenotypes and fate of effector PbTIIs using scRNA-seq. Our previous high-resolution scRNA-seq study suggested (Lonnberg et al., 2017), after high-dose transfer of 10^6^ PbTIIs, and assessment of tens to hundreds of PbTIIs, that two main effector fates, Th1 and Tfh were generated. Here, we transferred a physiologically relevant number of PbTIIs (10^4^ cells, which seeded the spleen with 100-500 cells) (Figure S4A), infected with *Pc*AS, and treated mice with sodium artesunate or control saline from peak (Figure 3A). At 7, 10, 14, 17, 21 and 28 days *p.i.* splenic PbTIIs were recovered by cell-sorting (Figure S4B), and processed via plate-based Smart-seq2. From 4548 wells, high quality data was obtained for 2964 (65.2%) cells (Figure 3A & S4C). Principle component analysis (PCA) revealed two main populations at day 7 *p.i.* (Figure 3B), which expressed either Th1 or Tfh gene signatures identified from our previous study (Figure 3B and Figure S5). To determine if a third rare subset might be present at peak, we performed an independent study of ∼11,000 PbTIIs at day 7 *p.i.*, compared to naïve PbTIIs using droplet-based Chromium scRNA-seq, powered to detect a rare third subpopulation if present at 1% frequency or more (Figure S6). PCA suggested two dominant populations from all day 7 cells, that exhibited clear CXCR5/6 bifurcation, and the same Th1/Tfh gene signature bifurcation as above (Figure S6A&B). Both Th1 and Tfh subsets also exhibited minor tails of highly proliferative, *Mki67-* expressing cells, which we inferred to be Th1/Tfh cells that had yet to complete clonal expansion (Figure S6B). Although *Ccr7* and *Sell* have been employed as markers for T_CM_ precursor cells, we saw no evidence for a population of T_CM_ precursors that could be readily distinguished transcriptomically amongst *Cxcr5-*expressing, Tfh-like cells (Figure S6B). Thus, taken together, these data suggested that PbTIIs gave rise to only Th1 and Tfh phenotypes.

**Figure 3.**
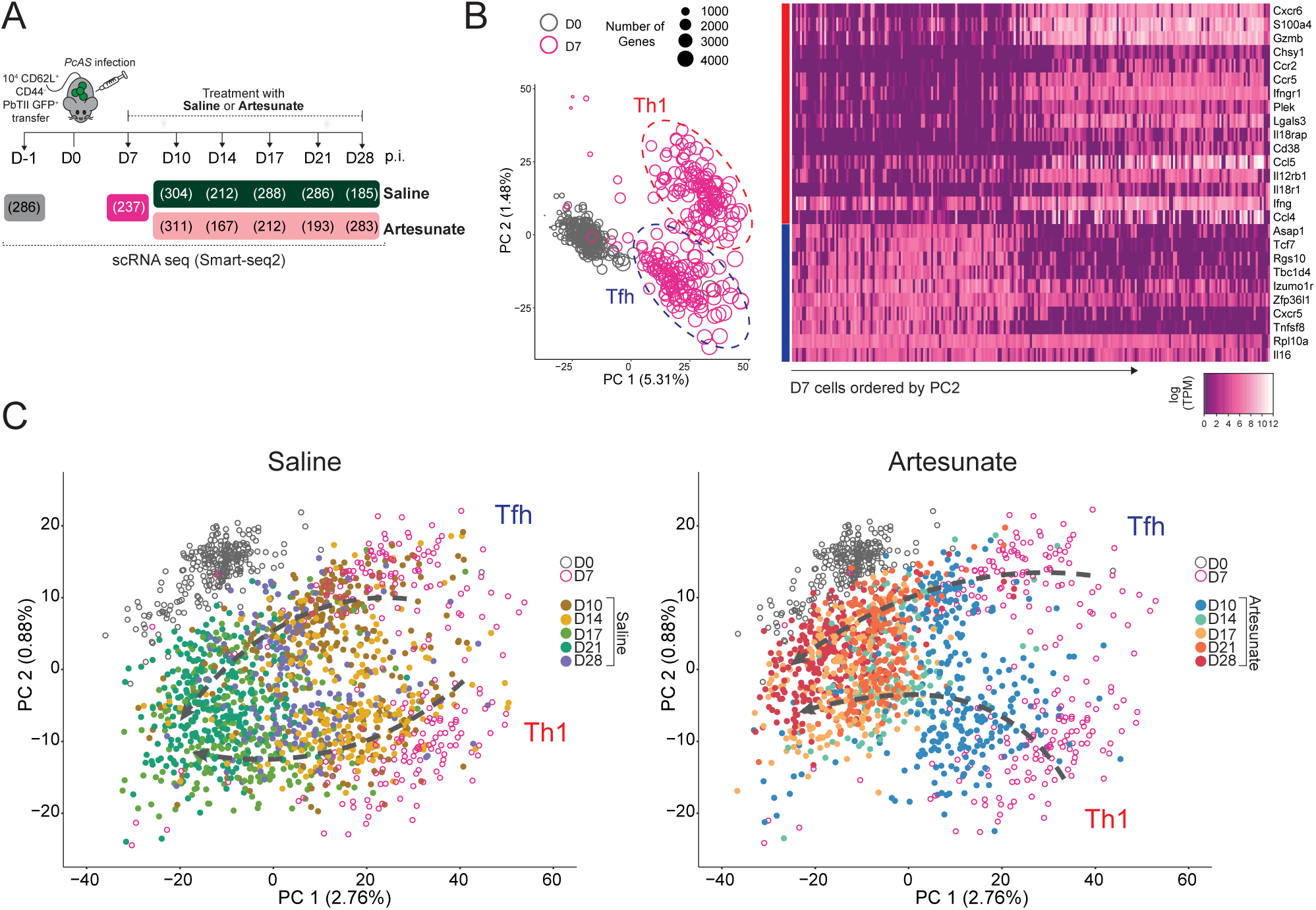
scRNA-seq assessment of memory PbTII cell development revealed two clusters at late timepoints. **(A)** Schematic of scRNA-seq experiment. PbTII cells were isolated at naïve state (D0, Figure S4A) and during effector to memory transition (D7-D28 *p.i.*, Figure S4B) for scRNA-seq assessment using the Smart-seq2 platform. Recipient mice (n=6/ group/timepoint) received 10^4^ sorted naïve (CD62L^+^ CD44^−^) GFP-expressing PbTII cells from a single donor 24 hours prior to infection with *PcAS*. At each timepoint, one mouse was chosen from each group for isolation of PbTII cells for scRNA-seq assessment. Numbers in parentheses denote the number of single cells that have passed quality check. **(B)** *(left)* PCA of PbTII cells from D0 and D7 *p.i.* The size of the circle represents the total number of genes expressed by each cell. *(right)* Heatmap describing expression of 16 Th1- and 10 Tfh-bifurcating genes derived from (Lonnberg et al., 2017), for each PbTII cell from D7 *p.i.*, ordered according to PC2 (*left*). **(C)** PCA of all PbTII cells as in (A). Analysis was performed on all cells and plotted as subsets of cells based on the treatment each mouse has received. Cells from D10 - D28 p.i. (closed circles) were either isolated from saline-(*left*) or artesunate-treated (*right*) mice. Cells isolated from D0 and D7 *p.i.* (open circles) were similarly replicated on both PCA plots. PCA was performed based on all genes expressed ≥100 TPM for >15 cells, with no regression for any variables (regression of variables such as cell cycle variation (Figure S7), mapped count variation (data not shown) revealed minimal effect on dataset).

We next conducted PCA on the Smart-seq2 time-course dataset (Figure 3C). Cells from one timepoint appeared adjacent and partially overlapping with previous and subsequent timepoints (Figure 3C). The existence of two cell states, marked by *Cxcr5* or *Cxcr6* expression, was evident at days 10 and 14 *p.i.* in saline and artesunate-treated mice (Figure 3C & S7A). This suggested progressive change in PbTII Th1 and Tfh transcriptomes over time, which was independent of cell-cycling effects (Figure S7B). By the third and fourth week, it was more difficult via transcriptomics to distinguish two clear sub-populations, although *Cxcr5* and *Cxcr6* were clearly expressed along one apparent branch but not the other (Figure S7A). It was also noted as parasitemia dropped (Figure S7C), and cells reverted to oxidative phosphorylation as a primary energy source (Figure S7D), that the number of detected genes dropped from ∼4000 at peak effector phase to less than 2000 towards memory (Figure S7E). Somewhat unexpectedly, memory PbTIIs expressed fewer genes than naïve PbTIIs (Figure S7E). Thus, PCA suggested possible trajectories taken by Th1 and Tfh cells after peak effector phase. The routes of these trajectories were altered by artesunate treatment, which appeared to nudge both trajectories closer towards naïve cells (Figure 3C). The data therefore suggested progressive change had occurred to the transcriptomes of Th1 and Tfh PbTIIs as they developed into memory cells, and that persisting infection had influenced the trajectories.

### Temporal mixture modelling reveals the molecular trajectories of Th1 and Tfh cells

Since Th1 and Tfh cells exhibited progressively altered transcriptomes during the second to fourth week of infection, we sought to reconstruct the entire transcriptomic response, from naivety through priming, clonal expansion, effector fate choice, contraction and memory. We combined Smart-seq2 data from this current study with two other PbTII datasets (Lonnberg et al., 2017), which had focused on the first week of infection. After batch correction (Figure S8), based on GPfates (Lonnberg et al., 2017) we performed Bayesian Gaussian Process Latent Variable Modelling (BGPLVM) to learn a minimum number of latent variables in the data. Two latent variables, LV1 and LV2, were sufficient to display much of the variability in the dataset (Figure 4A). BGPLVM of the combined datasets once again demonstrated overlapping cells between adjacent timepoints, and as in our previous study. We next generated a probabilistic model, firstly by calculating pseudotime coordinates for all cells based on LV1 (Figure S9A-B), and secondly, by running Overlapping Mixtures of Gaussian Processes (OMGP) analysis on saline or artesunate data separately, including data from day 0-7 in both datasets (Figure 4A). OMGP analysis revealed fate bifurcation within day 4 cells and two separate fates around day 7 *p.i.* (Figure 4A-B). Examination of Th1 and Tfh gene signatures from our previous study confirmed that the trajectory closer to naïve cells was Tfh-like, while the pathway tracking further away from naïve cells was the Th1-lineage (Figure S10). After peak effector phase, both for saline and artesunate groups, the ability to assign cells to either one lineage or the other began to diminish over pseudotime (Figure 4B). This was more apparent for saline-treated mice, where the two lineages mapped by OMGP converged substantially (Figure 4B). In contrast, Th1 and Tfh lineages remained distinct for longer after artesunate treatment (Figure 4B). Thus, the OMGP models provided high-resolution transcriptome dynamics and lineage tracking for PbTIIs for the first four weeks of infection. Numbers of detected genes increased from ∼2000 in naïve cells to ∼5000 during clonal expansion, which reduced to ∼4000 at effector phase, and progressively further to <2000 by the end of pseudotime, consistent with changes observed in cell cycle activity (Figure 4C-D). Anti-malarial drug treatment from effector phase did not alter the rate of decline in numbers of genes detected over pseudotime (Figure 4C-D), but as expected, did reduce the expression of T cell exhaustion-associated genes (Figure 4E). These data are consistent with the artesunate OMGP model describing effector to memory transitions in PbTIIs, while the saline OMGP model may be a more complex mixture of exhaustion and memory. Of interest, memory PbTIIs isolated from artesunate-treated group were observed to be positioned slightly later along pseudotime compared to saline-treated memory PbTIIs (Figure S9C & S11). Taken together, analysis of single PbTIIs over four weeks uncovered transcriptomic pathways between effector cells and their memory counterparts, revealing that effector to memory transitions are characterized not by rapid change, but by progressive modification to effector phenotypes.

**Figure 4.**
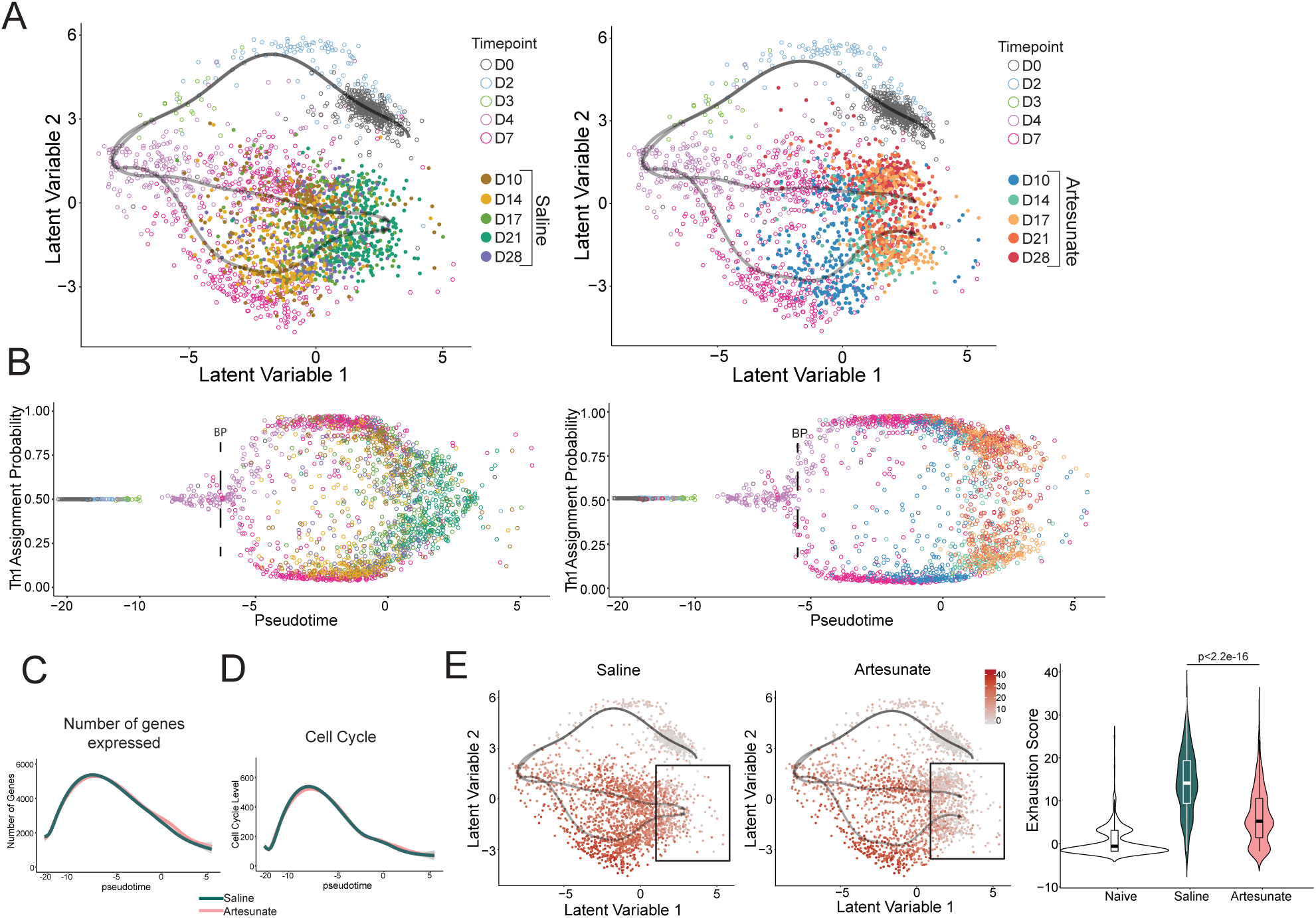
Temporal mixture modelling for tracing memory PbTII cell development during infection. **(A)** Reconstruction of trajectories along pseudotime by OMGP (Figure S8-S9). BGPLVM analysis was performed on the full time series from the combined PbTII datasets (Figure S8; 1^st^ batch: Smart-seq2 (384) D0-D28 *p.i.*, 2^nd^ batch: SMARTer C1 D0-D7 *p.i.*; 3^rd^ batch: Smart-Seq2 (96) D0, D4, D7 p.i.), and 2D representation was plotted based on treatment groups (left=saline; right=artesunate). D0, D2, D3, D4 and D7 *p.i.* cells were similarly replicated on each plot. OMGP analysis was performed separately for saline and artesunate. **(B-E)** “Saline” and “Artesunate” groups are comprised of the combination of shared timepoints (D0-D7) and unique timepoints based on treatment groups (D10-D28, saline/artesunate) from here on. **(B)** Th1 assignment probability assigned using OMGP along pseudotime for cells as described in (A). **(C)** Histogram of changes in number of genes expressed along pseudotime. **(D)** Histogram of cell cycle score along pseudotime. Each cell was scored for their cell cycle level based on the summative expression of 226 cell cycle genes derived from Cyclebase (Santos et al., 2015). **(E)** (*left*) Degree of exhaustion visualised on 2D representation of BGPLVM. Each cell was scored for their exhaustion phenotype based on the summative expression of 8 co-inhibitory receptors (*Ctla4, Pdcd1, Lag3, Havcr2, Btla4, Cd160, 2b4, Tigit*), established for their role in T cell exhaustion (Wherry and Kurachi, 2015). (*right*) Violin plots of exhaustion score from naïve cells (D0), and late cells positioned at late pseudotime (pseudotime>0.9) for saline and artesunate treatment groups (shown in black window). Statistical analysis was performed using Wilcoxon-rank sum test.

### Persisting infection differentially alters the transcriptomes of Th1 versus Tfh-derived memory PbTIIs

Given that the developmental trajectories taken by PbTII Th1 and Tfh cells were modified by drug treatment, we next determined transcriptional differences triggered in memory cells between saline and artesunate groups. We confined our analysis to single-cell transcriptomes positioned at the end of pseudotime, for either the Th1 or Tfh branch (Figure S12A). We performed differential gene expression analysis of “late pseudotime” Th1 or Tfh-lineage cells, for saline versus artesunate (Figure S12A). Firstly, we found the Th1 lineage was more sensitive to persisting infection than the Tfh lineage, since artesunate treatment altered the expression of 1110 genes (FDR < 0.05) in the Th1 lineage (Table S1) compared to 252 genes (FDR < 0.05) for the Tfh lineage (Table S2). Gene Ontology (GO) analysis revealed that in general, artesunate treatment increased the expression of immune-associated genes in the Th1 pathway, but had a more balanced effect on immune genes in the Tfh pathway, suggesting differing effects of persisting infection on each pathway. 175 immune genes were upregulated in the Th1 pathway after artesunate treatment, of which 17 were also upregulated in the Tfh pathway, including *Il7r, Cd96* and *Cd40lg.* (Figure S12B). Conversely, 31 immune genes were downregulated in the Tfh pathway, of which 20 were also downregulated in the Th1 pathway, including *Ctla4, Lag3, Il10, Tigit, Icos, Il21, Cd3d/g*. Effects on protein expression of Tigit, Lag-3 and IL-7R were subsequently all confirmed by flow cytometry (Figure S12C-E). Together, our data suggested that although persisting infection triggered a partial exhaustion phenotype in both lineages, immune function genes were more adversely affected late in the Th1 pathway than the Tfh pathway.

### Th1 and Tfh lineage tracing reveals developmental relationships with other CD4^+^ T cell subsets

OMGP analysis of Th1 and Tfh effector PbTIIs revealed two transcriptomic pathways leading towards memory/exhaustion. These pathways permitted examination of the extent to which effector phase gene expression was retained long term. *Cxcr5* and *Cxcr6* displayed strong expression with slight waning along each respective trajectory (Figure 5A). These were better at distinguishing between lineages than master transcription factors, *Tbx21* or *Bcl6*, or *Ifng* (Figure S13A). Of note, *Tbx21* and *Ifng,* though Th1-defining in many instances, were expressed here in both lineages, illustrating Th1-characteristics were evident throughout the entire dataset after bifurcation. This was further supported by expression patterns for Th1-associated chemokine receptor gene, *Cxcr3* (Figure S15A).

**Figure 5.**
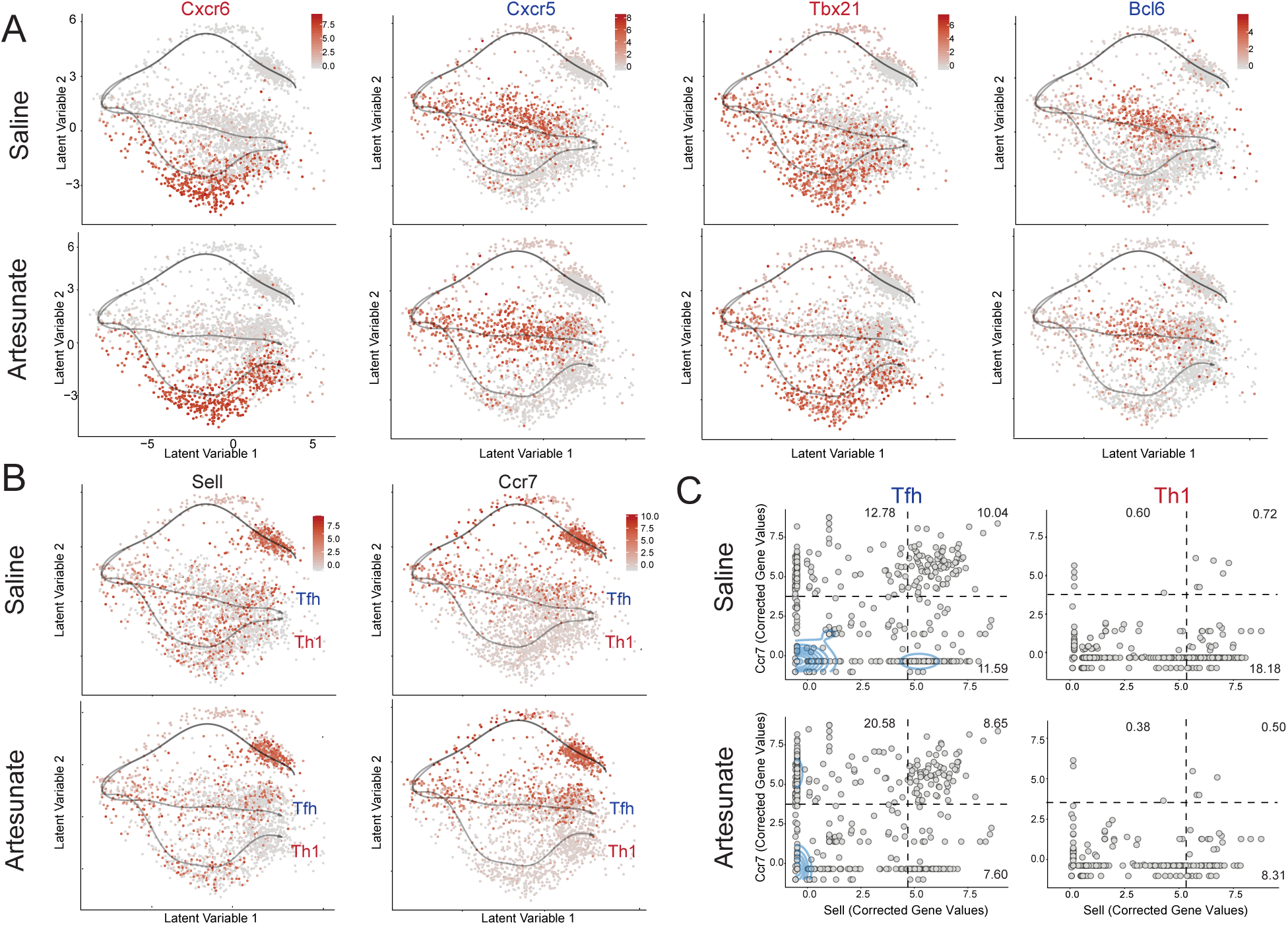
Th1 and Tfh memory PbTII cells are developmentally linked to other CD4^+^ T cell subsets. **(A)** Visualisation of Th1-(*Cxcr6, Tbx21*) and Tfh-(*Cxcr5, Bcl6*) associated genes on 2D BGPLVM representation. **(B)** Visualisation of *Sell* and *Ccr7* on 2D BGPLVM representation overlaid with OMGP trajectories representing Th1 or Tfh branch. **(C)** Co-expression of *Sell* and *Ccr7* along pseudotime, for all cells annotated for the Tfh (left) or Th1 (right) branch as outlined in methods (Figure S10C). Kernel density estimation failed to estimate a population of *Sell^+^ Ccr7^+^* cells, hence a threshold was estimated where cells express both *Sell* ≥ 4.7 and *Ccr7* ≥ 3.75 corrected gene values.

Two classical markers of T_CM_ cells are CCR7 and CD62L, which are also expressed at high levels by naïve CD4^+^ T cells. We noted that although *Sell* (encoding CD62L) was uniformly downregulated by PbTIIs immediately after priming (Figure 5B), *Ccr7* was retained by a subset of PbTIIs during clonal expansion, and was subsequently retained by a subset of cells specifically along the Tfh lineage (Figure 5B). Thus, while *Sell* appeared to be stochastically expressed over time and between lineages, *Ccr7* was restricted to the Tfh lineage. ∼10% of cells in the Tfh lineage co-expressed *Sell* and *Ccr7,* while co-expression was almost absent in the Th1 lineage (Figure 5C). We also considered whether GC Tfh cells were detectable in the Tfh lineage by assessing *Pdcd1* expression (which encodes PD1). *Pdcd1* and *Ccr7* were stochastically expressed within the Tfh lineage (Figure S14A), but with relatively little overlap (Figure S14B). This suggested that while *Pdcd1^+^ Cxcr5^+^* GC Tfh cells and *Ccr7^+^ Cxcr5^+^* emerging T_CM_ cells might be present in the Tfh-pathway, our dataset had insufficient power to differentiate between them. Nevertheless, these data are consistent with PbTIIs along the Tfh lineage giving rise to either or both GC Tfh cells, T_CM_ and memory Tfh cells.

We next considered fates along the Th1 lineage. Our previous study reported emerging co-expression of *Il10* and *Ifng* amongst PbTIIs, consistent with their classification as Tr1 cells (Lonnberg et al., 2017). Here, we noted *Ifng* expression, though not restricted to the Th1 lineage, was nevertheless higher along this pathway compared to the Tfh lineage (Figure S13A). *Il10* expression was heavily restricted to the Th1 lineage, with 35% of Th1 pathway cells co-expressing these two cytokines (Figure S13B). Anti-malarial drug treatment reduced the incidence of Tr1 cells towards the end of pseudotime (Figure S13B), consistent with a requirement for persisting infection to promote their retention. Together, these data indicate that the majority of Tr1 cells developed progressively from Th1 effectors in this infection model, and that their capacity to express *Il10* relied on persisting infection. Expression of *Ifng,* however, was less affected by drug treatment, consistent with PbTIIs being poised as T_EM_ cells for Th1 recall. Therefore, our analysis is consistent with Th1 cells giving rise directly to Tr1 cells, which progressively give rise to Th1-poised T_EM_ cells. In summary, our pathway inference analysis suggested that Tfh cells ultimately give rise to T_CM_ cells, memory Tfh and/or GC Tfh, while Th1 cells progress via a Tr1 intermediate state to give rise to Th1-poised T_EM_ cells.

### Clonality analysis via endogenous TCR chains reveals varying predisposition to Th1/Tfh fate choice

In our previous study, we had used heterogeneous endogenous TCR chain expression in *Rag1^+/+^* TCR transgenic PbTIIs as molecular barcodes (Lonnberg et al., 2017). We identified six families (2-3 cells/family), each derived from one naïve PbTII cell, and noted three instances where single naïve PbTIIs had given rise to both Th1 and Tfh fates. Here, we reasoned that transfer of 100-fold fewer cells would reduce inter-clonal competition, and increase the chances of detecting clonally related cells. Consistent with this, from 2964 Smart-seq2 transcriptomes across all timepoints and conditions, we reconstructed endogenous VDJ regions using TraCeR (Stubbington et al., 2016), and detected 201 families (Figure 6A & B), defined by cells sharing the same endogenous alpha and beta chain sequences. Family sizes ranged from 2-5 cells (Figure 6C). As expected, family members did not cross timepoints or conditions (data not shown), because individual mice were used for each, and PbTIIs transferred to each mouse were expected to have unique endogenous barcodes. This indicated the fidelity of our endogenous barcoding approach. Next, consistent with our previous study, we found it common, though not universally so, for cells from the same family to exhibit both Th1 and Tfh fates (Figure 6B). This revealed that individual naïve PbTIIs often gave rise to progeny that progressed along both the Th1 and Tfh pathways. Thus, even in this model where functional TCR diversity was absent, single naïve CD4^+^ T cells displayed heterogeneity in the fates adopted by their progeny. We next examined whether Th1/Tfh cell fate choice was influenced by cell family. Examining only cells in artesunate-treated mice, since distinct binary fates were more easily discerned across all time points compared to saline controls (Figure 6B), and excluding cells where fate was indeterminate, we focused on 97 families. We reasoned that if a cell’s fate was independent of its family, the distribution of cell fates within each family should follow a random, binomial distribution (Figure 6D). However, we observed that PbTIIs of the same family were more likely to commit to the same fate than predicted by a random binomial process (p<0.0005, Figure 6D). Therefore, adoption of either fate by the progeny of a single naïve PbTII cell was not random. Rather, our mathematical modelling suggested a variable propensity between clonotypes for the progeny of a single naïve CD4^+^ T cell to progress along either a Th1 or Tfh pathway towards memory. Given that all cells expressed the same functional TCR, these data are consistent with T cell-extrinsic factors differentially controlling the fate of cells derived from the same progenitor.

**Figure 6.**
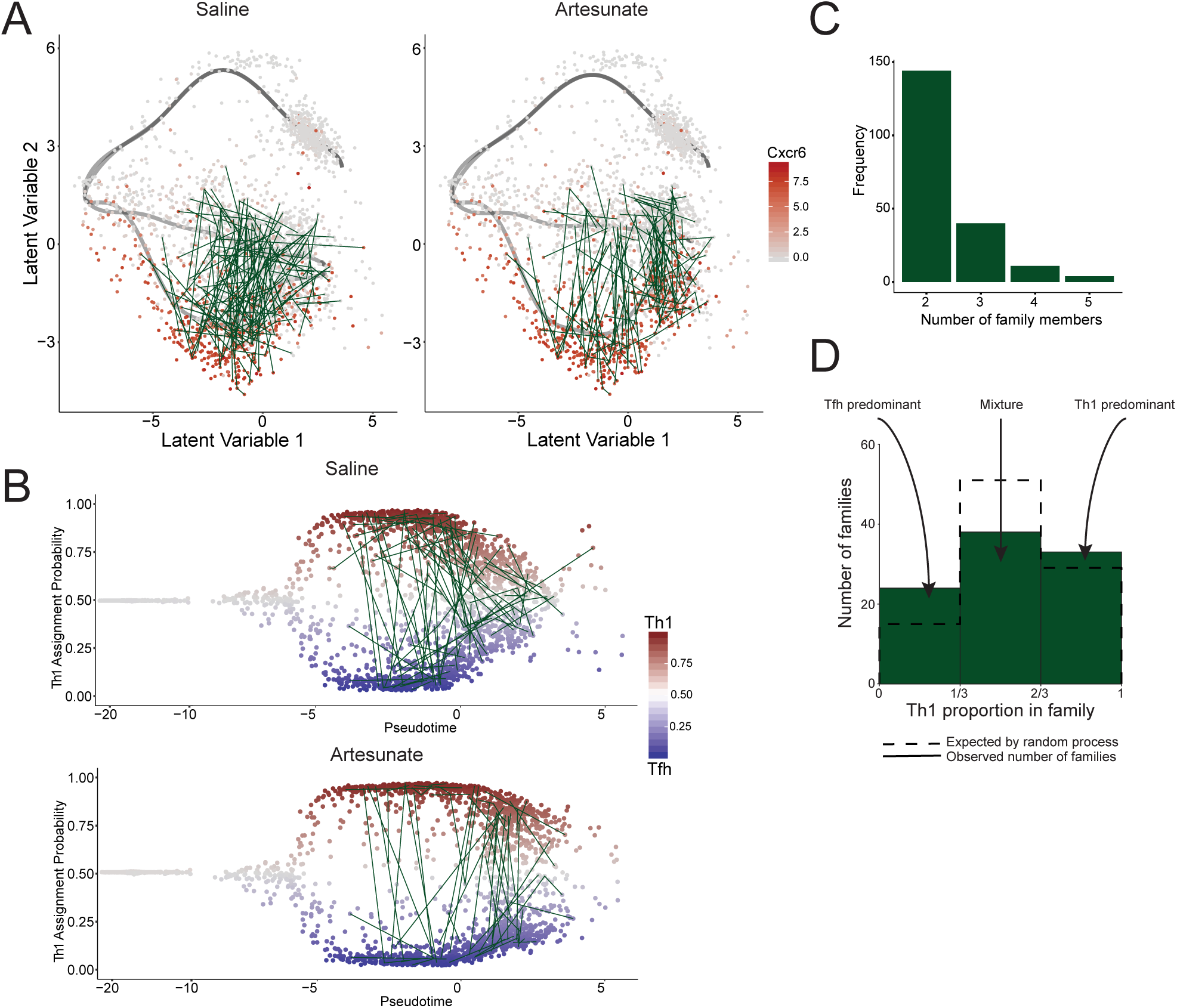
The relationship between T cell clonotypes and Th1/Tfh fate bifurcation. **(A)** Visualisation of families sharing endogenous TCR α and β sequences on 2D BGPLVM representation in saline-treated group (124 families) and artesunate-treated group (97 families), with each family connected by green lines. Cells are coloured by their *Cxcr6* expression. Families were seen to occupy both Th1 and Tfh fates along pseudotime. **(B)** Visualisation of families on Th1 assignment probability along pseudotime. **(C)** Bar graph representing number of families for each family size. **(D)** Bar graph representing observed number of families (solid green bar) that are (from left to right): Tfh predominant, mixture of Tfh and Th1, and Th1 predominant. Dotted empty bar represents expected number of families by random process as described in methods.

### Pseudotime correlation analysis reveals a gene signature for memory CD4^+^ T cells

We next reasoned that progressive increases in gene expression along pseudotime, along either or both lineages, would reveal those genes associated with memory onset. Individual genes, for both the saline or artesunate OMGP models, were given two scores, firstly for the strength of their association with either Th1 or Tfh lineage, and secondly for the degree to which their expression correlated with pseudotime (Figure 7A). The vast majority of genes correlated with neither pseudotime nor Th1/Tfh, and remained neutral in the assessment. In contrast, a small number of genes exhibited strong Th1 or Tfh assignment scores and correlated moderately with pseudotime (Figure 7B & C). These included *Cxcr6, Ccr2* and *Ccr5* for Th1-associated genes (Figure 7C). For those genes with Th1-bias, we noted several that correlated with late pseudotime, including *Ccl5*, *Nkg7*, *S100a4* and *S100a6* (Figure 7C). In contrast, few if any Tfh-skewed genes correlated with late pseudotime. Instead, many Tfh-lineage associated genes correlated with very early pseudotime values, revealing that Tfh-associated genes are also expressed at high levels in naïve or recently activated CD4^+^ T cells (Figure 7A). Finally, we noted a subset of genes exhibiting little Th1/Tfh correlation, but whose expression correlated with very late pseudotime. We reasoned these genes, often similarly observed in saline and artesunate groups, were memory-associated (Figure 7A, 7D-E). Of interest were transcription factors *Id2* and *Maf*, whose roles in CD4^+^ T cell memory remain unclear. We confirmed elevated expression of ID2 and the reciprocally related protein, TCF1 (encoded by *Tcf7*) (Masson et al., 2013), in a subset of PbTIIs at day 28 *p.i.* (Figure S15D, E & G). Interestingly, we also noted differential expression of *Id2* and *Id3*, with *Id2* being expressed in both lineages and *Id3* restricted to the Tfh pathway (Figure S15D & F). *Cxcr3* expression also correlated highly with late pseudotime, suggesting this marker though conventionally associated with Th1 cells, may be a general marker for Th1-conditioned memory in *Plasmodium* infection (Figure 7A-B). Consistent with this, we detected cell-surface CXCR3 protein on most PbTIIs during persisting infection at day 28 *p.i.* (Figure S15B), as well as on CXCR6^+^ and CXCR5^+^ PbTIIs after artesunate treatment (Figure S15C). Thus, our pseudotime correlation analysis identified a memory-associated gene signature particularly after removal of persisting infection, which unlike proliferative genes expressed tightly during pseudotime (Figure 7E), remained strongly upregulated after four weeks of infection and anti-malarial drug treatment (Figure 7E).

**Figure 7.**
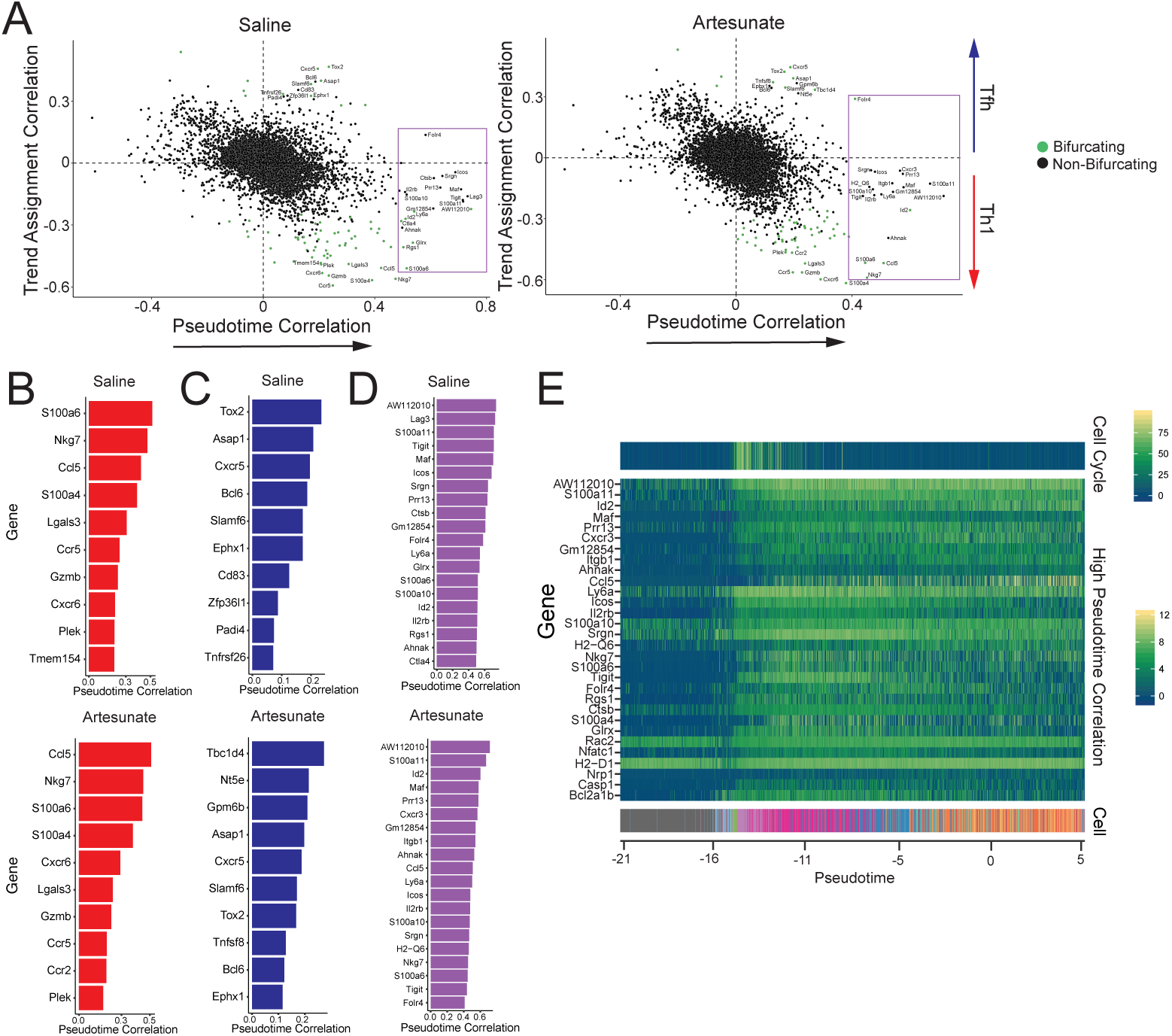
Changes in gene dynamics accompanying CD4^+^ T cell memory differentiation. **(A)** Identification of genes associated with pseudotime progression. Gene expression correlation with pseudotime (x-axis) versus the correlation with Th trend assignment (y-axis) is shown (Pearson). Genes were calculated for their relevance with Th bifurcation using GPfates (denoted in green), for each dataset separately. Top 20 pseudotime correlated genes are labelled and enclosed within the purple window. **(B)** Ranking of top 10 Th1-bifurcating genes based on trend assignment probability according to their correlation with pseudotime. **(C)** Ranking of top 10 Tfh-bifurcating genes based on trend assignment probability according to their correlation with pseudotime. **(D)** Ranking of genes based on their correlation with pseudotime as identified in (A). **(E)** Expression patterns of top 30 pseudotime-correlated genes along pseudotime for cells from the artesunate-treated group. Sum of 20 cell cycle gene expression is shown at the top.

To validate our memory-associated gene signature, we performed an independent scRNA-seq experimental repeat, and compared the transcriptomes of naïve PbTIIs with those from day 28 *p.i.*, with and without sodium artesunate from peak. Using droplet-based scRNA-seq on the Chromium platform and subsequent PCA (Figure S16A), we again observed memory PbTIIs that were transcriptomically more heterogeneous than naïve cells (Figure S17A), and that removal of persisting infection brought PbTII transcriptomes slightly closer to naïve than saline controls (Figure S16A). We observed a retention of the *Cxcr5*/*Cxcr6* bifurcated structure under both conditions with a tail of proliferative cells in saline controls (Figure S16B&C), as well as evidence of cells with enriched *Ccr7* and *Sell* expression in the *Cxcr5^+^* cluster (Figure S16C). Most importantly, the gene signature identified from OMGP modeling of Smart-seq2 data also differentiated between naïve and day 28 *p.i.* cells (Figure S16D). Therefore, in using two separate scRNA-seq platforms and temporal mixture modelling, we identified a gene signature associated with the acquisition of memory.

### scATAC-seq reveals loss of heterogeneity and partial resetting of chromatin during memory onset

scRNA-seq had suggested that PbTIIs following the Th1 and Tfh trajectories became progressively similar to each other and closer to naïve cells during memory onset. Since these pathways were generally associated with an onset of transcriptional quiescence, we hypothesized that apparent partial convergence of lineages was an artefact of transcriptomic assessment. To test this, we examined the epigenomic phenotype of PbTII cells using single-cell Assay for Transposase Accessible Chromatin by sequencing (scATAC-seq). Specifically, we tested whether loss in transcriptomic and protein marker heterogeneity during effector to memory transition was evident at chromatin level. We assessed PbTIIs at naïve (day 0), expansion (day 4), peak (day 7) and memory/exhaustion (day 32 *p.i.* with or without persisting infection) (Figure 8A). To aid Th1/Tfh classification of individual epigenomes, we also cell-sorted CXCR5^hi^ and CXCR6^hi^ PbTIIs at day 7 *p.i.* (Figure 8A). Across the entire scATAC-seq dataset, we identified 59,197 peaks of accessible chromatin. PCA of all accessible peaks indicated greatest variability along PC1 was attributed to the number of reads sequenced, consistent with previous observations (Chen et al., 2018). PC2 separated recently activated PbTIIs (day 4 & 7 *p.i.*) from naïve and day 32 *p.i.* cells, while PC3 separated CXCR6^hi^ from CXCR5^hi^ PbTIIs within day 7 *p.i.* (Figure 8A). Thus chromatin accessibility in PbTII effectors was substantially more heterogeneous than in naïve PbTIIs. Importantly, epigenomes of memory or exhausted PbTIIs clustered closer to naïve PbTIIs than to effectors, and were more homogeneous than effectors (Figure 8A). Targeted analysis of Th1/Tfh associated binding motifs showed differential and inverse patterns of accessibility for *Id2* and *Tcf7* in CXCR6^hi^ (Th1) cells compared to CXCR5^hi^ (Tfh) cells, that was partially reset to naïve levels by day 32 *p.i.* (Figure 8B). *Tbx21* motifs were enriched for, and *Bcl6* motifs depleted from peaks during the first week of infection, which again was partially lost by day 32 *p.i.* (Figure 8B). These data revealed expected changes in chromatin accessibility for specific Th1/Tfh-associated transcription factors during the first week of infection, which was partially retained in memory. More broadly, 2700 (46.4% of total) accessible peaks in naïve PbTIIs were present in distal enhancer regions, while only 371 (30.4%) distal enhancer peaks were detected in memory PbTIIs generated in the absence of persisting infection. Reactome pathway analysis revealed an enrichment for peaks associated with tyrosine-kinase and CD28-signalling genes in memory PbTIIs, that was absent in naïve PbTIIs (Figure 8C). Together these data suggested that despite partial resetting of chromatin from effector phase, specific changes in chromatin landscape had been retained in memory cells.

**Figure 8.**
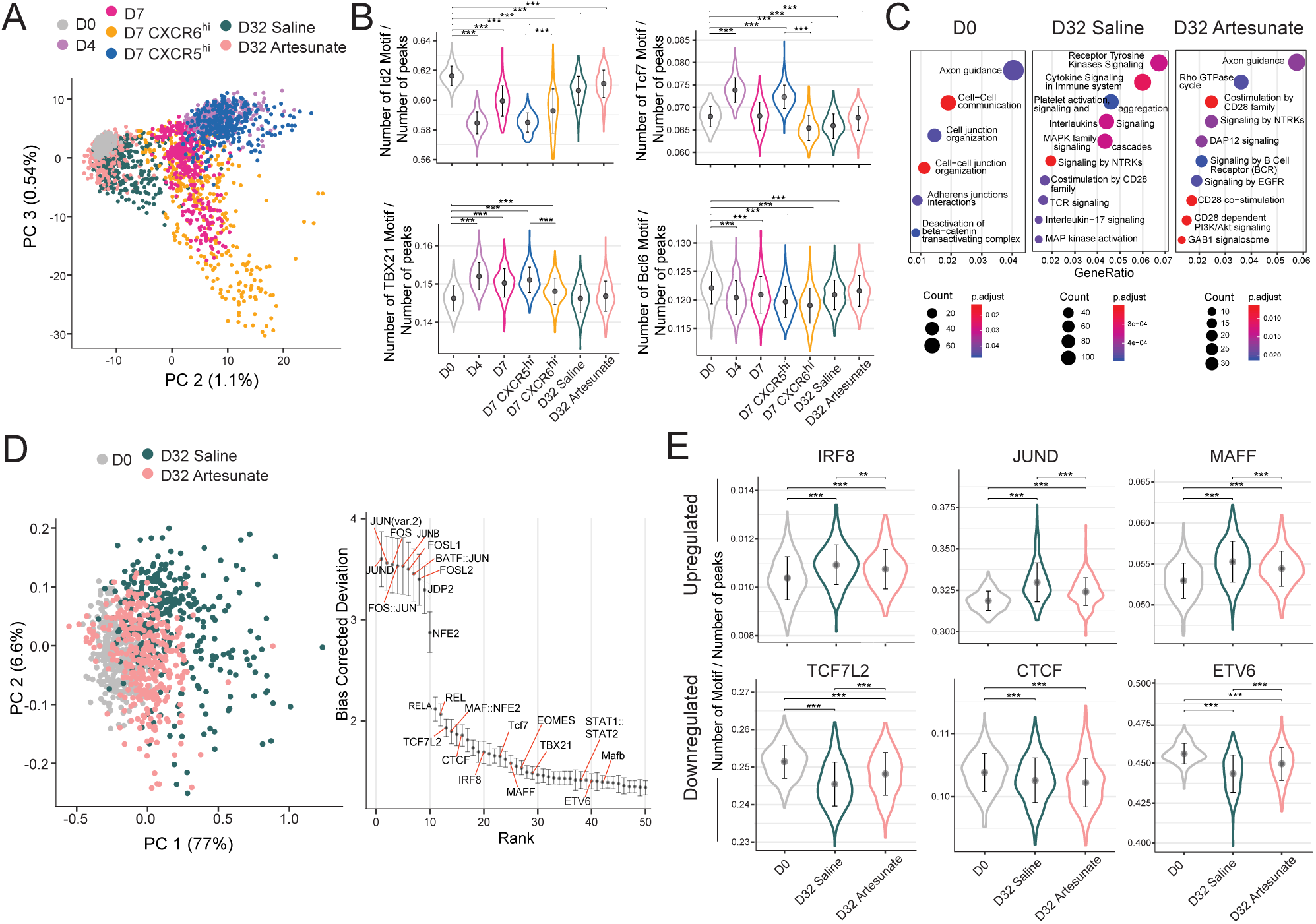
scATAC-seq assessment of changes in chromatin landscape during memory PbTII development. **(A)** PCA of peaks after application of Latent Semantic analysis (LSA) (Cusanovich et al., 2018). Binarised peaks as “open” or “closed” from all cells were merged for LSA application. **(B)** Changes in fractions of accessible chromatin regions associated with different Th1 or Tfh-associated transcription factor *(Id2, Tcf7, Tbx21, Bcl6*) for different timepoints. Two-sided t-test was performed for all combinations, but only relevant statistical results were displayed. **(C)** Reactome analysis of top 10 enriched pathways for different timepoints. P-value was adjusted for multiple testing using Benjamini-Hochberg correction. The size of the circle represents the number of genes from our dataset that were involved in each individual pathway. The ratio for the number of genes from our dataset involved in each pathway, from the overall total number of genes known for each of those pathways were calculated and denoted as “gene ratio”. **(D)** (*left*) PCA of 514 variable motifs (combined database of human and mice motifs from JASPAR database) calculated using chromVAR analysis for comparison between naïve (D0), saline and artesunate-treated cells from D32 *p.i.* (*right*) Top 50 motifs explaining variability between naïve (D0), saline and artesunate-treated cells from D32 *p.i.* For each motif, a deviation value is calculated for each cell representing how different the accessibility for peaks with that motif is from the expectation based on all cells being equal, with further correction for biases. **(E)** Changes in fractions of accessible chromatin regions associated with different transcription factors detected from chromVAR analysis as described in (D).

Next, we sought to identify new transcriptional regulators whose motif accessibility varied between naïve and memory cells, as an indication of possible biological relevance (Schep et al., 2017). PCA of motif bias-corrected deviation revealed that day 32 *p.i.* motif variability was similar though distinct from naïve PbTIIs (Figure 8D). Binding motifs for several transcription factors were variable including those from the Fos, Jun, Runt, NF-kappaB, Maf, Tbx, Tcf7 and Irf families (Figure 8D and Table S3). We detected binding motif enrichment for IRF8, JUND and MAFF, and depletion of TCF7, CTCF, ETV6, NFATC3, and RUNX1 motifs in memory cells relative to naïve cells (Figure 8E & Figure S17). Taken together, our data indicated that genome-wide heterogeneity in chromatin accessibility in effector PbTIIs was partially, but not completely, reset during memory transition.

## Discussion

Previous fate mapping, adoptive transfer and single-cell tracking studies in acute viral and bacterial infection support a model in which memory CD4^+^ T cells develop directly from effector precursors (Hale et al., 2013; Harrington et al., 2008; Marshall et al., 2011; Pepper et al., 2011; Tubo et al., 2016). However, the existence of molecular pathways governing these transitions has remained unclear, due to difficulties in detecting transient, intermediate cellular states, and a reliance upon small subsets of marker proteins. Here, we attempted to overcome these challenges by using high resolution single-cell transcriptomics combined with computational modelling, in an experimental model of non-lethal malaria. Our model controlled for the effect of TCR diversity on effector fate choice by using *Plasmodium-*specific TCR transgenic CD4^+^ T cells. Importantly, PbTIIs primed within the same *Pc*AS-infected recipient generated similar proportions of both Th1 and Tfh effectors, which allowed us to examine the fate of, and potential relationships between these effector sub-types in the same system.

We hypothesized that progressive transcriptomic change occurs in those Th1 and Tfh cells that give rise to memory CD4^+^ T cells. We captured and re-ordered single-cell transcriptomes not according to their time-point of capture, but according to their position along an as yet undefined developmental trajectory. In doing so, we revealed two developmental lineages originating from Th1 and Tfh-like effector states. These lineages both exhibited a gradual reduction in the numbers of genes expressed, from ∼4000 genes at effector phase to less than 2000 at memory. It was notable that naïve counterparts expressed 2000 genes, suggesting that memory CD4^+^ T cells may transcribe a more limited set of genes than naïve cells, although this remains to be tested.

Various models have been proposed to describe the relationship between effector and memory T cells, including divergent, linear and branching models. Aside from the divergent model, in which naïve T cells give rise early to separate effector and memory lineages, most other models imply that effector cells, perhaps at different stages of their development, give rise to memory cells. Our study captured intermediate states between effector and memory, and provides evidence that supports two linear developmental pathways. Firstly, Th1 cells gave rise directly to *Il10/Ifng* co-expressing Tr1 cells, which subsequently gave rise to quiescent cells expressing higher levels of *Tbx21, Ifng, Cxcr6,* and less *Ccr7* than either memory cells in the other lineage or naïve cells, consistent with being T_EM_ cells poised for Th1-recall responses. Secondly, the Tfh lineage ultimately gave rise to quiescent cells expressing higher levels of *Ccr7, Cxcr5, Tcf7 and Bcl6* compared to T_EM_ cells, consistent with them being T_CM_ cells. While microscopy and flow-cytometry confirmed the existence of PD1^hi^ CXCR5^+^ GC Tfh amongst splenic PbTIIs, it was not possible to clearly discern separate lineages for T_CM_ versus GC Tfh, or indeed memory Tfh cells. While a recent scRNA-seq dataset from LCMV-infection suggested separate populations of Tfh and T_CM_ precursors were detectable, our data using both Smart-seq2 and the Chromium platform found little supporting evidence for transcriptomically separate Tfh and T_CM_ precursor populations, but some suggestion in the Chromium dataset of separate memory Tfh and T_CM_ cells within the Tfh pathway. Based on these data, we speculate that early Tfh cells and T_CM_ precursors are in fact the same cell-type in the PbTII/*Pc*AS model. While some early Tfh-like PbTIIs may progress to memory Tfh cells, and others towards *Ccr7*-expressing T_CM_ cells, their transcriptomic developmental pathways could not be readily dissected via transcriptomics in this study. This might suggest that Tfh and T_CM_ precursors are in fact different terms for the same transcriptomic state, or perhaps that our study was not powered sufficiently to determine the subtle differences between them.

While our data revealed a correlation between memory onset and *Id2* upregulation along both lineages, we also noted only along the Tfh lineage re-expression of *Tcf7* and its encoded protein, TCF-1, after being lost from naïve cells. *Tcf7* has been reported to promote Tfh cell differentiation as well as CD4^+^ T cell development in the thymus, but has yet to be fully examined for roles in controlling CD4^+^ T cell memory. Given that *Tcf7* may control self-renewal capacity in T cells (Lin et al., 2016; Nish et al., 2017), we hypothesize that *Tcf7* is important for promoting the persistence of T_CM_ cells and/or memory Tfh cells, but not T_EM_ cells.

Consistent with CD4^+^ T cell responses in bacterial and viral infection models, splenic PbTII numbers contracted from peak by >95% over the subsequent three weeks. While our single-cell transcriptomic approach captured those cells which progressed to memory along two lineages, we were not able to readily detect apoptotic or pre-apoptotic PbTIIs during contraction from day 7-14 *p.i..* Given that spleen cells went through multiple processing stresses, including mechanical tissue homogenization, exposure to hypotonic buffers to lyse red blood cells, and turbulence and shear forces during cell-sorting, it is possible that fragile pre-apoptotic cells were simply lost in the procedure. We speculate that less disruptive approaches, such as spatial transcriptomics (Rodriques et al., 2019), may help determine why some PbTIIs progress to memory, while the majority do not.

PCA and OMGP analysis revealed that although both pathways tended to return towards naïve cells, the Tfh-T_CM_ pathway remained closer to naïve PbTIIs compared to the Th1-Tr1-T_EM_ lineage, even at the end of the month. This is consistent with a view that like naïve CD4^+^ T cells, T_CM_ cells exhibit a renewing capacity, and are not poised to produce effector cytokines as readily as T_EM_. Interestingly, curtailing persisting infection, which removed the confounding factor of T cell exhaustion, increased expression of GO terms and genes associated with basic cellular integrity in T_CM_ cells, and increased expression of GO terms and immune function-related genes in T_EM_ cells. Moreover, persisting infection had a more disruptive effect on T_EM_ cell biology compared to more quiescent T_CM_ cells. Our transcriptomic model (Figure 9) therefore indicates that Tfh-T_CM_ transitions remain much closer to naïve T cell biology, and are more resistant to T cell exhaustion compared to those progressing along the Th1-Tr1-T_EM_ lineage.

**Figure 9.**
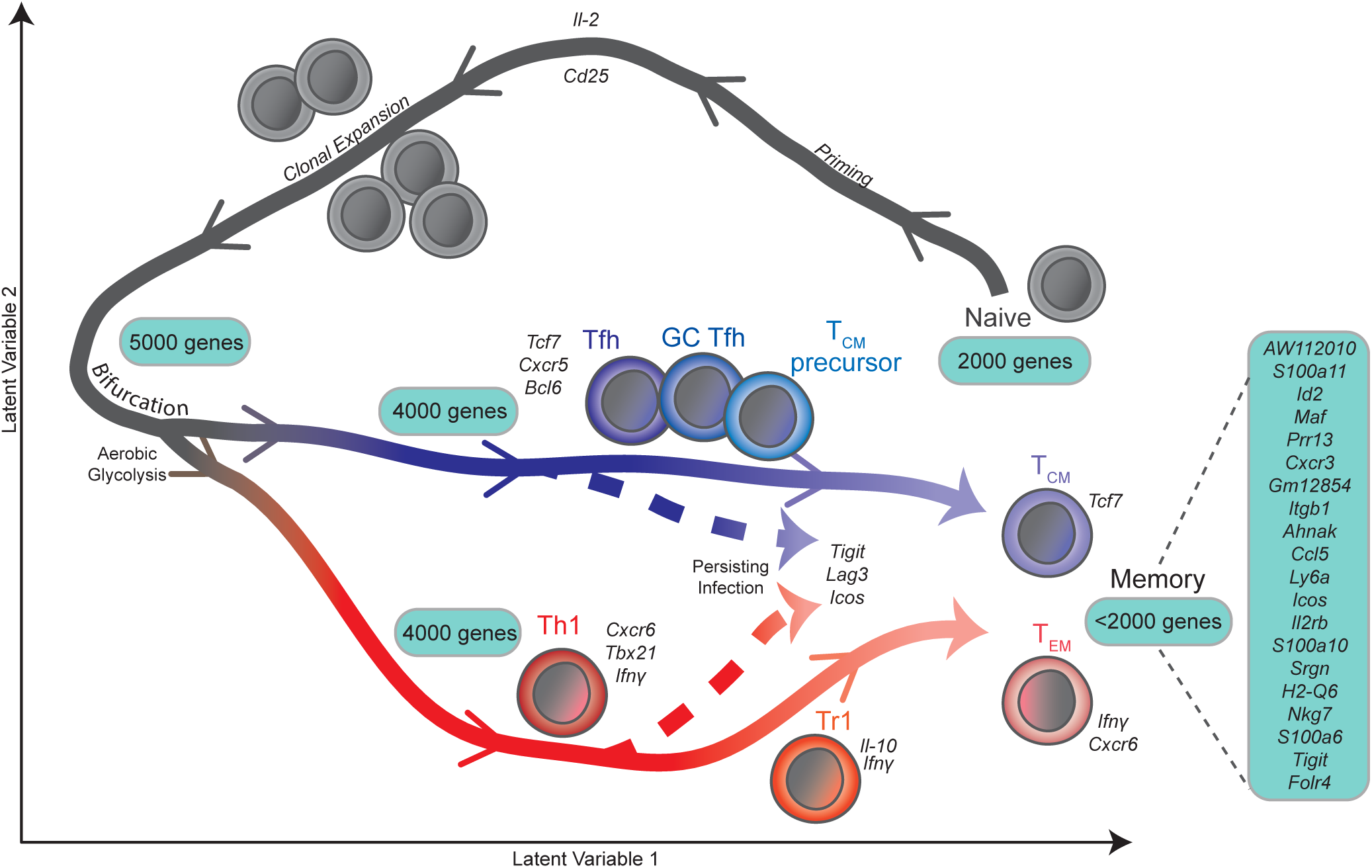
A conceptual view for CD4^+^ T cell memory development. A broad overview of the transcriptomic dynamics accompanying transition of CD4^+^ T cells from naivety to memory. CD4^+^ T cells appear to branch into effector Th1 and Tfh after undergoing clonal expansion, and exhibit a partial retention of similar effector Th fates as they transition into memory cells. Th1 effector cells give rise to Th1-poised T_EM_ memory cells with Tr1 as an intermediary subset. On the other hand, Tfh effector cells give rise to T_CM_ cells with shared characteristics with GC Tfh and/ or memory Tfh cells. The presence of a persisting infection subtly diverted development of memory CD4^+^ T cells (dashed lines) by increasing the expression of exhaustion-associated signature. Detection of a number of genes that correlated with memory development are summarised in the box (right). Numbers in boxes denote the average total number of genes expressed at different stages of CD4^+^ T cell differentiation.

By using TraCeR on high-resolution, full-length transcript scRNA-seq data, to examine low-level transcription of diverse, endogenous TCR sequences in *Rag-*sufficient PbTIIs, we detected 201 clonotypes, or families, of PbTIIs, each derived from an individual naïve PbTII cell. This number represented a substantial increase from our previous study (Lonnberg et al., 2017), and provided an opportunity to assess the frequency with which cells from the same family adopt the same or different fates. Firstly, consistent with previous reports, single naïve PbTIIs were capable of giving rise to more than one fate, confirming that diversity in effector fate is common amongst clones. Most importantly, fitting to a mathematical model indicated that random acquisition of Th1 or Tfh fate for cells in any given family was unlikely for ∼50% of all families. These gave rise in some families a strong predisposition to either one fate or the other. This suggested heterogeneity amongst clonal cells, which raises the question of what promotes this effect. Although PbTIIs were not strictly clonal due to being on a *Rag-*sufficient background, we previously showed this played no role in controlling Th1/Tfh fate choice (Lonnberg et al., 2017). Instead, we noted substantial heterogeneity in the micro-anatomical location of PbTIIs at peak, relative to T and B cell zones. Given that B cells and myeloid cells control PbTII effector fate (Fernandez-Ruiz et al., 2017; James et al., 2018; Lonnberg et al., 2017) we hypothesize that cell-cell interactions are primary drivers of memory fate. Assuming that progeny from a naïve CD4^+^ T cell stay within the same vicinity after clonal expansion, at least before and during effector differentiation, it is possible that similar T cell-extrinsic signals and cell-cell interactions in the area will promote the same effector and memory fate amongst a family. We noted that our model was not sensitive to the timepoint of capture, suggesting that memory cells of the same family had been subjected to the same influences as effectors. Our analysis also suggest that for ∼50% of cell families, acquisition of Th1 or Tfh fate was more random within the family. Many different non-mutually exclusive scenarios might account for this alternative outcome including greater cell motility negating the influence of micro-environments within the spleen, asymmetric cell division, and internal stochasticity. Thus, our data suggest that multiple mechanisms likely contribute to memory fate choice within and between families, and that the balance between T_EM_ and T_CM_ proportions is ultimately dependent upon the balance of Th1/Tfh fate choices within individual families and populations as a whole. We speculate that micro-anatomical location and velocity of activated CD4^+^ T cells are the primary determinants of effector and memory fate choice for any given antigen-specificity.

Temporal mixture modelling of single-cell RNA-seq data permitted the discovery of genes whose expression kinetics correlated with memory onset. We defined a small subset of genes associated with memory onset in both lineages, including the transcription factors *Id2* and *Maf,* chemokine-related genes, *Ccl5* and *Cxcr3,* integrin *Itgb1*, and cytokine receptor *Il2rb*. We hypothesize this gene set may be useful for distinguishing naïve from memory states in polyclonal CD4^+^ T cell populations, particularly after Th1-biased immune-challenges. It was notable that *Cxcr3*, which is upregulated early, before PbTIIs bifurcated along Th1 or Tfh fates, was also strongly retained at the end of the month. It has been noted in mouse models of malaria, and in humans infected with *Plasmodium*, that Tfh cells often acquire Th1 characteristics including CXCR3 expression (Obeng-Adjei et al., 2015; Ryg-Cornejo et al., 2016). The existence of CXCR5^+^ and CXCR6^+^ memory populations that co-expressed CXCR3 suggest that Th1-like Tfh memory cells or Th1-like T_CM_ may also be generated in this model.

An important feature of the two lineages was their progressive, partial coalescence during memory onset. This was apparent at both transcriptomic and epigenomic levels, substantiating our view that memory development involves a decrease in heterogeneity amongst effector cells, and a return to chromatin structures that to a certain extent resemble naïve CD4^+^ T cells. Importantly, however, changes in accessibility to transcription factor binding motifs were preserved amongst individual memory compared to naïve cells. For example, MAFF and IRF8 motifs were retained in memory cells relative to naïve; given the increased transcription of *Maf,* and *Irf8* (data not shown), during memory onset, we hypothesize these transcription factors may play specific roles in CD4^+^ T cell memory.

Transcriptomic dynamics revealed that low-level persisting infection subtly impacted the pathways towards memory, but did not completely prevent effector-to-memory transition. Curtailing infection triggered an upregulation in immune function genes in the Th1-Tr1-T_EM_ lineage, consistent with improved recall capacity, while genes associated with basic cellular function and integrity were upregulated along the Tfh-T_CM_ lineage. These data suggest that persisting infection had differing effects on the two lineages, although the precise effects on memory Tfh and/or T_CM_ cells remains to be determined. Our study provides support for linear models of development for CD4^+^ T cell memory, where Th1 cells give rise to T_EM_ cells, and Tfh cells give rise to memory Tfh and/or T_CM_ cells. It will be important to examine whether similar developmental pathways exist in other models in which, for example, Th2 and Th17 effector cells or tissue resident memory CD4^+^ T cells are generated.

In summary, our study indicated that linear transitions from effector to memory occur in a gradual, progressive manner. Our data were less supportive of a branched model, in which memory precursors develop during the effector response. The extended timeframe over which transcriptomic change occurred to effector CD4^+^ T cells suggests potential opportunities for manipulating memory development after a primary immune response has been elicited.

## Methods

### Experimental mice, adoptive transfer and infections

C57BL/6J mice were purchased from Animal Resources Centre (Canning Vale), PbTII (Fernandez-Ruiz et al., 2017) and nzEGFP (Bhat et al., 2014) mice were bred in-house. nzEGFP mice express enhanced green fluorescent protein (EGFP) from the ubiquitously expressed CMV early enhancer/chicken beta actin promotor. All mice were female and aged 6-12 weeks and were maintained under specific pathogen-free conditions within the animal facility at QIMR Berghofer Medical Research Institute. All animal procedures and protocols were approved (A1503-601M) and monitored by the QIMR Berghofer Medical Research Institute Animal Ethics Committee.

Spleens were collected and homogenised through a 100 µm cell strainer to create a single cell suspension. Red blood cells (RBC) were lysed using RBC Lysing Buffer Hybri-Max™ (Sigma-Aldrich) and CD4^+^ T PbTII cells were enriched using CD4 microbeads (Miltenyi Biotec). Cells (10^4^/200 µl) were transferred to each mouse via lateral tail vein intravenous (i.v.) injection.

*PcAS* parasites were used after thawing frozen, infected blood stabilites and performing a single *in vivo* passage in C57BL/6J mice. *PcAS-*infected RBCs were harvested from passage mice by cardiac puncture and mice infected with 10^5^ *PcAs* parasites via lateral tail vein i.v. injection.

### Treatment with anti-malarial sodium artesunate

Sodium artesunate (Guilin Pharmaceutical Co., Ltd.) was prepared according to the manufacturer’s protocol by diluting in 0.9% saline (Baxter) to a final concentration of 5 mg/ml. Mice were treated via intraperitoneal (i.p) injection with 200 µl sodium artesunate (1 mg/mouse), or vehicle control saline, twice daily from day seven to day nine post-infection (*p.i.*), once daily from day 10 to day 16 *p.i.* and then twice weekly until experimental endpoint. Mice treated with sodium artesunate were also administered pyrimethamine (0.07g/L; Sigma-Aldrich) in drinking water for the duration of the treatment.

### Parasitemia assessment

Parasitemia assessment of mice infected with *PcAS* was carried out as previously described (James et al., 2018; Lonnberg et al., 2017). Briefly, a single drop of blood was collected via tail bleed and diluted in 250 µl of RPMI medium containing 5 U/ml heparin sulphate. Diluted blood was stained with Syto84 (5 µM; Life Technologies) and Hoechst 33342 (10 µg/ml; Sigma-Aldrich) for 30 minutes in the dark at RT. Staining was quenched with 10 volumes of ice-cold RPMI medium and samples immediately acquired by flow cytometry.

### Flow cytometry

Spleens were collected and homogenised through a 100 µm cell strainer to create a single cell suspension and RBCs were lysed using RBC Lysing Buffer Hybri-Max™ (Sigma-Aldrich). Splenocytes were assessed for viability using a LIVE/DEAD™ Fixable Aqua Dead Cell Stain Kit (Life Technologies), according to the manufacturer’s protocol, unless otherwise specified. Prior to antibody staining, Fc receptors were blocked using antibodies against CD16 and CD32. Cells were incubated with surface marker antibodies (Table 1) for 20 minutes at 4°C. To assess cytokine’s and transcription factor expression, cells were incubated with brefeldin-A (10 mg/ml) with or without ionomycin (500 ng/ml) and PMA (25 ng/ml) at 37°C for 3 hours. Intracellular staining was then performed using the eBioscience™ Foxp3/Transcription Factor Staining Buffer Set. Samples were acquired on a LSRII Fortessa analyser (BD Biosciences) and subsequently analysed using FlowJo software (Treestar).

### *In vivo* re-challenge

C57BL/6J mice received enriched naïve, GFP-expressing CD4^+^ T PbTII cells (10^4^ cells/ mouse) 24 hours prior to infection with *PcAS* parasites (10^5^ pRBCs/ mouse). Mice were treated with artesunate or saline as vehicle control according to dosing regimen described. Treatment with artesunate or saline ceased at 23 days *p.i.* Mice then received a new batch of enriched naïve, congenically-marked (CD45.1^+^) CD4^+^ PbTII cells (10^5^ cells/ mouse) at day 27 *p.i.*, prior to re-challenge with 10^7^ *PcAS* pRBCs 24 hours later.

### Immunofluorescence microscopy

Spleens were fixed with 2% paraformaldehyde for 2-4 hours at RT in the dark followed by dehydration with 30% sucrose (Chem-supply) overnight at RT in the dark. Spleens were snap-frozen in Tissue-Tek Optimal Cutting Temperature embedding media (Sakura Finetek) on dry ice and stored at −80°C. Spleens were sectioned at 10-30 μm on Polysine slides such that consecutive sectioning is avoided. Sections were allowed to dry overnight. The dried sections were re-hydrated for 15-20 minutes before fixation in 4% paraformaldehyde for 15-20 minutes at RT in dark. Slides were washed three times, for 5 minutes in washing buffer (0.01% Tween20 in PBS) before permeabilisation with 0.1% Triton-X in washing buffer for 10-15 minutes. After washing, slides were incubated with Medical Background Sniper (Biocare) for 30 minutes. Slides were rinsed for 2 minutes and endogenous biotin was blocked using an Avidin/Biotin Blocking kit (Vector Laboratories, Inc.), according to manufacturer’s protocol. Tissue sections were then stained with rabbit anti-GFP (Abcam), rat anti-mouse CD3-AF594 (Biolegend), rat anti-mouse IgD-AF647 (Biolegend), and biotinylated peanut agglutinin (PNA; Vector Laboratories, Inc.) for 1-2 hours at RT in the dark. Secondary antibody staining for PNA and GFP was performed using streptavidin-AF555 (Thermo Fisher Scientific) and donkey anti-rabbit AF488 (Thermo Fisher Scientific), respectively, for 1-2 hours at RT in the dark. Tissue sections were incubated with DAPI for 10-15 minutes to counterstain nuclei, and slides were mounted in Dako Mounting Media (Agilent Technologies). Image acquisition was performed using an Aperio FL slide scanner or a Zeiss 780-NLO point scanning confocal microscope at 20x, 40x and 63x objective.

Cell detection and quantification was performed using the spot detection function in Imaris image analysis software (Bitplane), with thresholds <10 μM. All objects were manually inspected for accuracy before data were plotted and analysed in GraphPad Prism. Co-localisation analysis was performed using Imaris co-localisation functions. B cell follicle was defined as IgD-positive region, while GC was defined as IgD-negative and PNA-positive region within B cell follicle.

### Bulk Fast-ATAC sequencing and analysis

Fast-ATAC sequencing was performed as previously described (Corces et al., 2016). Briefly, 5,000 viable PbTII cells were sorted by flow cytometry and pelleted by centrifugation. Supernatant was removed and 50 µl of transposase mixture (25 µl TD buffer (Illumina), 2.5 µl of TDE1 (Illumina), 0.5 µl of 1% digitonin (Promega), 22 µl nuclease-free water) added to the cells. Cells were then incubated at 37°C for 30 min at 300 rpm in an Eppendorf ThermoMixer. Transposed DNA was amplified and purified using a QIAGEN MinElute PCR Purification kit, according to the manufacturer’s protocol. Purified DNA was eluted in 20 µl Buffer EB (QIAGEN) and sequenced using paired-end sequencing on a NextSeq 550 instrument (Illumina).

Raw ATAC-seq reads were mapped to mouse genome MGSCv37 (mm9) using BWA-MEM (Li, 2013). Unmapped reads, reads mapping to unassigned contigs and mate unmapped reads were removed, as well as mitochondrial genes and PCR duplicates. The resulting bam files were first converted to bedGraph format using the bedtools “genomecov” command from the BEDTools suite (Quinlan and Hall, 2010) and then converted to bigwig format using the bedGraphToBigWig program from University of California, Santa Cruz (UCSC) Genome Browser (https://genome.ucsc.edu/). Read counts were normalised to the number of uniquely mapped reads per million. Peak calling was performed using MACS2 (Zhang et al., 2008) with the following parameters: --nomodel, --shift 37, --extsize 73, --pvalue 1e-5. The bigWig tracks and narrowPeak files were visualised in a custom UCSC Genome Browser track hosting the mouse mm9 reference genome. Peaks overlapping with the mm9 blacklist region generated by UCSC were removed and overlapping peaks between replicate samples were identified using the “findOverlaps” command from the GenomicRanges package (Lawrence et al., 2013) and used for downstream analysis. Peaks were annotated using the R package ChIPseeker (Yu et al., 2015) with transcription start site (TSS) reaching from −3000 to 3000 to the UCSC mm9 gene model and the “org.Mm.eg.db” R package.

### Single-cell RNA capture and sequencing

#### Smart-seq2

384-well lo-bind plates were prepared with 0.5 µl of Triton-X lysis buffer, 0.25 µl of 10 µM oligo-dT30-VN, 0.25 µl dNTP mix (25 mM each) and ERCC controls (final dilution of 1:64 million) per well and stored at −20°C until use. Plates were thawed on ice before sorting single PbTII cells into each well. Single-cell lysates were sealed and spun at 100 × *g* for 1 minute then immediately frozen on dry ice and stored at −80°C. Reverse transcription and PCR were performed following the Smart-seq2 protocol (Picelli et al., 2014) with the following modifications: 1 µl of reverse transcription mix and 5.7 µl PCR Master mix. After amplification, cDNA was subjected to quality control using 1 µl of amplified cDNA on an Agilent 2100 BioAnalyser and Agilent High Sensitivity DNA kits (Agilent Technologies). 5 µl of cDNA per cell was cleaned using Agencourt AMPure XP beads (Beckman Coulter) at a 1.0x ratio on a Hamilton STAR liquid handler (Hamilton Robotics). cDNA was quantified using Biotium AccuClear High Sensitivity DNA quantification reagent and normalised to 1 ng/µl in a total volume of 500 nl before library preparation using Nextera XT DNA Sample Preparation Kit (Illumina). 125 nl of in-house index adapters (Integrated DNA Technologies), similar to Illumina N7 and N5 indices, were added to the tagmentation reaction before adding 1.5 µl of KAPA HiFi DNA polymerase (KAPA Biosystems) and performing 12 cycles of PCR according to the manufacturer’s instructions. After PCR, all samples were pooled into 384-plex pools using a 24×16 dual indexing approach and the pool cleaned using Agencourt AMPure XP beads at a 0.6x ratio. Library pools were eluted in buffer EB and quality controlled using an Agilent 2100 BioAnalyser and Agilent High Sensitivity DNA kits before adjusting the concentration to 4nM. Each library was sequenced on one lane of Illumina HiSeq 2000 version 4 chemistry (paired-end 75-base pair reads).

#### Chromium 10x Genomics

PbTII cells were isolated by flow cytometry into a 1% BSA/PBS buffer. Approximately 8,000 cells were loaded per channel onto a Chromium controller (10x Genomics) for generation of gel bead-in-emulsions. Sequencing libraries were prepared using Single Cell 3’ Reagent Kits v3 (10x Genomics) and then converted using the MGIEasy Universal Library Conversion Kit (BGI) before sequencing on a MGISEQ-2000 instrument (BGI).

### Processing of scRNA-seq data

#### Smart-seq2 data

Raw reads were mapped to a mouse genomic index (Ensembl version 70) using STAR aligner version 2.5.2 (Dobin et al., 2013) and Transcripts Per Millions (TPM) were estimated by RSEM version 1.2.30 (Li and Dewey, 2011).

#### 10x Genomics data

For the BGI FASTQ files to be made compatible with “cellranger count” pipeline from Cell Ranger version 3.0.2 (10x Genomics), file names and FASTQ headers were reformatted using code from https://github.com/IMB-Computational-Genomics-Lab/BGIvsIllumina_scRNASeq (Senabouth et al., 2019). Data was processed using 10x mouse genome 1.2.0 release as a reference.

### Quality control of scRNA-seq data

#### Smartseq2 data

Cells were filtered to remove those with less than 100,000 reads that mapped to the mouse genome, those expressing less than 1,000 genes and more than 5,000 genes, and those with more than 35% mitochondrial content. Only genes expressed at 1 or more TPM in 3 or more cells were considered, unless otherwise specified.

#### Chromium 10x Genomics data

Cells were filtered to remove those expressing less than 200 genes and more than 6,000 genes, and those with more than 35% mitochondrial content. Only genes expressed in 3 or more cells were considered.

### Batch effect correction

Two PbTII datasets from our previous study underwent quality control as described previously (Lonnberg et al., 2017) and were combined with Smart-seq2 data from our current study. Genes expressed at less than 1 TPM in less than 3 cells were globally removed. The “removeBatchEffect” function from Limma (Ritchie et al., 2015) was then used to remove the batch effect.

### Dimensionality reduction

PCA dimensionality reduction (prcomp) was performed on Smart-seq2 data with genes expressed at 100 or more TPM in 15 or more cells as input. On the combined, batch-effect corrected dataset, dimensionality reduction was performed with Bayesian Gaussian Processes Latent Variable Modelling (BGPLVM) using GPfates as previously described (Lonnberg et al., 2017).

To perform PCA dimensionality reduction on 10x Genomics data, highly variable genes were identified using “FindVariableGenes” function from Seurat (Butler et al., 2018) and were used as an input.

### Pseudotime inference and lineage tracing

Pseudotime was inferred on the combined, batch-effect corrected dataset based on latent variable (LV) 1 coordinates from BGPLVM dimensionality reduction. Firstly, DBSCAN clustering was performed on D0-D3 cells and D4-D28 cells separately using LV1 and LV2 coordinates as inputs (Ester et al., 1996). From D0-D3 group, only cells with LV2 coordinates lower than the lowest LV2 coordinate from the non-outlier cells were considered as outliers, and from D4-D28 group, only cells with LV2 coordinates higher than the highest LV2 coordinate from the non-outlier cells were considered as outliers. Outliers from each group were grouped together with the cells from the opposing groups. LV1 coordinates of the first group (D0-D3 cells + outliers from D4-D28 cells) were flipped along the lowest LV1 coordinate from all dataset and concatenated with the LV1 coordinates of the second group (D4-D28 cells + outliers from D0-D3 cells) to infer the pseudotime coordinates.

Lineages were traced along the pseudotime with Overlapping Mixtures of Gaussian Processes (OMGP) using GPfates (Lonnberg et al., 2017), using LV2 coordinates to model as a function of pseudotime. Lineage tracings were performed separately for saline and artesunate groups, with the same set of parameters used (two trends assumed, global variance = 0.5, per-trend variance = 3, and per-trend lengthscale = 2). Here, the saline group consists of all cells from D0-D7 and saline-treated cells from D10 onwards. The Artesunate group consists of all cells from D0-D7 and artesunate-treated cells from D10 onwards.

### Differential gene expression analysis

As an input for differential gene expression analysis (DGEA), expected counts from Smart-seq2 data was estimated by RSEM version 1.2.30 and then normalized and log-transformed using “computeSumFactors” and “normalize” functions from scran (Lun et al., 2016). Pairwise Welch t-test was performed using “pairwiseTTests” function from scran to identify genes differentially expressed between two groups of cells. Comparisons were done as follows: (1) cells at late pseudotime from artesunate group on the Th1-arm versus cells at late pseudotime from saline group on the Th1-arm, (2) cells at late pseudotime from artesunate group on the Tfh-arm versus cells at late pseudotime from saline group on the Tfh-arm (Figure 4E). Genes with false discovery rate (FDR) below 0.05 were considered as differentially expressed.

### Gene ontology term enrichment analysis

Gene ontology (GO) terms were obtained from “org.Mm.eg.db” Bioconductor annotation package (Carlson, 2019). Fisher’s exact test was used to identify significantly over-represented GO terms. Input gene lists were differentially expressed genes from comparisons described above, with up- and downregulated genes considered separately. The background gene set was reduced to those used as an input for DGEA. Enrichment analysis for biological process terms was performed using “goana” function in edgeR (Robinson et al., 2010).

### Testing the distribution of cell fates among clonotypes

T cell receptor (TCR) sequences for each cell from Smart-seq2 data was reconstructed using TraCeR as previously described (Stubbington et al., 2016). Cells were classified as Th1 group if their Th1 assignment probability (TAP) was greater than the median TAP + 0.05 and cells were classified as Tfh group if their TAP was less than the median TAP – 0.05.

Focusing our analysis on only those cells from the 97 families found in artesunate treated animals, and excluding cells where fate was indeterminate, we wished to test whether the distribution of cell fate amongst families was consistent with cells acquiring their fates independently of the family to which they belong, or if cell fate was linked to family. To test the null hypothesis that cells fate is randomly distributed across cells, independent of their family (i.e. ancestry), we used a bootstrapping approach. First, we constructed a likelihood function to describe the likelihood of observing the a given number of Th1 cells, *Th*_*i*_, in a particular family, *i*, of size *n*_*i*_, assuming the process is binomial with some probability *P*_*h*_. The overall likelihood of observing the distribution of cell fates across all families is then given by,

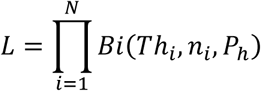

where *N* is the number of families. Note that the *P*_*h*_ that maximises the likelihood will be given by the proportion of all cells that have a Th1 fate (demonstrated by evaluating the derivative of when log of the likelihood function and solving for when derivative is zero). In this data set we observe that the proportion of cells that were Th1 was *P*_*h*_ = 0.575. Using the observed numbers of *Th*_*i*_ and *n*_*i*_ for each family, we compute L in equation 1 and call this the likelihood of the observed distribution (*L*_0_).

To determine whether the likelihood of the observed distribution of cell fates by family is consistent with a random process, cells were randomly permuted amongst the different families 100,000 times. Each permutation, *p*, was performed in such a way as to preserve the size of families, but redistribute the cells across the families. After permuting the cells, the likelihood, *L*_*p*_, for was recalculated for each permutation *p*. This process produced a distribution of the likelihood associated with cell fates being randomly and independently distributed across families. We then compare the likelihood of the observed distribution of cell fates across families, *L*_0_, with the set of likelihoods, *L*_*p*_, associated with each random permutation of cells across the fates and found that the likelihood of finding the observed distribution of cell fates by family by random chance was less than approximately 0.05%, i.e. *p* = 0.0005, thus allowing us to reject the null hypothesis that cell fate is independent of family.

The histogram of the observed versus expected distribution of families with a Th1 predominance, a Tfh predominance or a mixture of fates (Figure 6D), was determined by first drawing a histogram of the observed proportions of Th1 cells in each family (noting that families of size 3 or 6 that had a proportion of 1/3 or 2/3 were included in the middle bin in the histogram). To determine the expected distribution of families, the above described permutation of cells between families was performed and the histogram redrawn each time. The expected distribution of families was created by taking the mean of the bin counts across these 100,000 random permutations.

### Single-cell ATAC sequencing and processing

#### Plate-based scATAC-seq

Plate-based scATAC-seq was performed as previously described (Chen et al., 2018). Briefly, 17,000-50,000 viable PbTII cells were sorted using flow cytrometry and pelleted by centrifugation. Supernatant was removed and cells were resuspended in 50 μl tagmentation mix (33 mM Tris-acetate, pH 7.8, 66 mM potassium acetate, 10 mM magnesium acetate, 16% dimethylformamide (DMF), 0.01% digitonin and 5 μl of Tn5 from the Nextera kit from Illumina, Cat. No. FC-121-1030). Cell/ tagmentation mixture was then incubated at 37°C, 800rpm on an Eppendorf thermomixer for 30 minutes. Tagmentation reaction was stopped by the addition of equal volume (50 μl) of stop buffer (10 mM Tris-HCl, pH 8.0, 20 mM EDTA, pH 8.0) followed by incubation in ice for 10 minutes. 150 uL of DPBS/ 0.5% BSA was added to the mixture and nuclei suspension was then transferred to a FACS tube. 0.0006% DAPI (in-house) was added for staining the nuclei prior to single-nuclei sort into 384-well plate (Eppendorf, Cat. No. 0030128508) containing 2 µl of 2X lysis buffer (100 mM Tris.HCl, pH 8.0, 100 mM NaCl, 40 µg/ml Proteinase K (New England BioLabs, P8107S), 0.4% SDS. Single-nuclei lysates were sealed and spun at 100 × g for 1 minute then immediately frozen on dry ice and stored at −80°C.

#### scATAC-seq processing

Plates were taken out from −80°C, thawed at room temperature and incubated at 65°C for 15 minutes. 2 µl of 10% tween-20, 1 µl of 10 µM of indexed primer mix and 5 µl of NEB Next-High Fidelity 2X PCR Master Mix were added to each well in that order. PCR was performed as 72°C 10minutes, 98°C 5 minutes, [98°C 10 seconds, 63°C 30 seconds, 72°C 20 seconds] × 18. All reactions were combined, and the pooled library was purified using a Qiagen minElute PCR purfiation column. Library concentration was determined by Agilent Bioanalyzer 2100 High Sensitivity kit. Each 384 pool was sequenced on one lane of Illumina HiSeq 2000.

### scATAC-seq data processing

Raw ATAC-seq reads were mapped to mouse genome MGSCv37 (mm9) using BWA-MEM (Li, 2013). Reads with mapping quality below 30 and duplicated reads marked with Picard (http://broadinstitute.github.io/picard/) were removed. For each timepoint, all bam files were aggregated and duplicated reads were marked again and removed. Peak calling was performed using MACS2 (Zhang et al., 2008) with parameters -q 0.01 --nomodel --shift −100 --extsize 200 for each aggregated timepoint. The union of aggregated timepoint peaks (q < 0.01) was generated using bedops (Neph et al., 2012). To generate the full read count matrix over the union of peaks for each single cell, bedtools “coverageBed” command was used (Quinlan and Hall, 2010).

#### Quality control metrics

For each cell, the numbers of total reads sequenced, uniquely mapped reads, duplicated reads and mitochondrial reads were calculated. In addition, the fraction of reads in peaks and the fraction of open chromatin were calculated as previously described (Chen et al., 2018).

#### Filtering cells and peaks

To identify low quality cells for each timepoint, cells were marked if log2-transformed uniquely mapped reads counts, mitochondrial content and fraction of open chromatin were twice as high as the median absolute deviation. If a cell deviates twice from any of the three metrics, it was removed prior to analysis. Peaks were removed if they overlapped with the mm9 blacklist region generated by UCSC. Peaks mapped to regions present in less than 30 cells across all cells were removed.

#### Dimensionality reduction

The filtered coverage matrix was binarised and used as input for latent semantic analysis (lsa) followed by principle component analysis using R package lsa (https://CRAN.R-project.org/package=lsa).

#### Motif in peaks fraction and motif variability

To assess the role of transcription factor (TF) motif accessibility during memory formation R packages Biostrings (Pagès, 2019), TFBSTools (Tan and Lenhard, 2016) and transcription factor motif database JASPAR2016 (Tan, 2019) were utilised. For each TF motif of interest, each peak was annotated if the underlying sequence contained the motif with a minimum matching score of 80%. Then for each cell, the number of peaks with the TF motif were counted. The number of peaks were then corrected with a motif by the number of total peaks of that cell. In addition, TF motif variability across all time points or selected time points of interest were assessed using chromVar (Schep et al., 2017) with default parameters. In total, 514 TF motifs were assessed from mouse and human from the JASPAR2016 TF motif database.

#### Peak annotation and Enrichment analysis

Peaks were annotated with R package ChIPseeker (Yu et al., 2015) with transcription start site (TSS) reaching from −3000 to 3000 to the UCSC mm9 gene model and the “org.Mm.eg.db” Bioconductor annotation package (Carlson, 2019). Genes with TSS closest to peaks that were identified solely during peak calling of the aggregate of D0, D32 Saline or D32 Artesunate were assessed for pathway enrichment using R package ReactomePA (Yu and He, 2016).

**Table 1.**
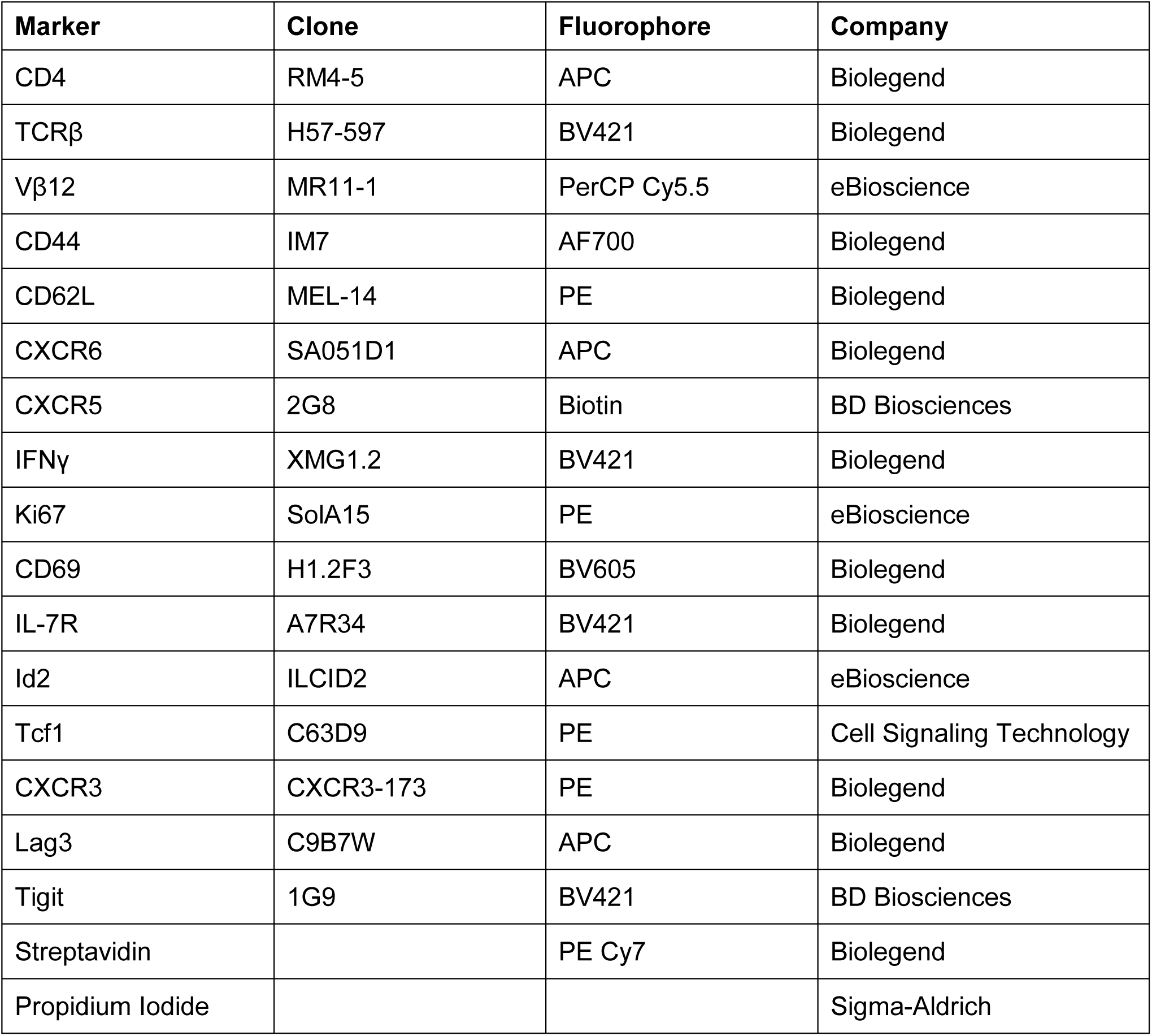
Flow cytometry antibodies.

---

**Supp 1.**
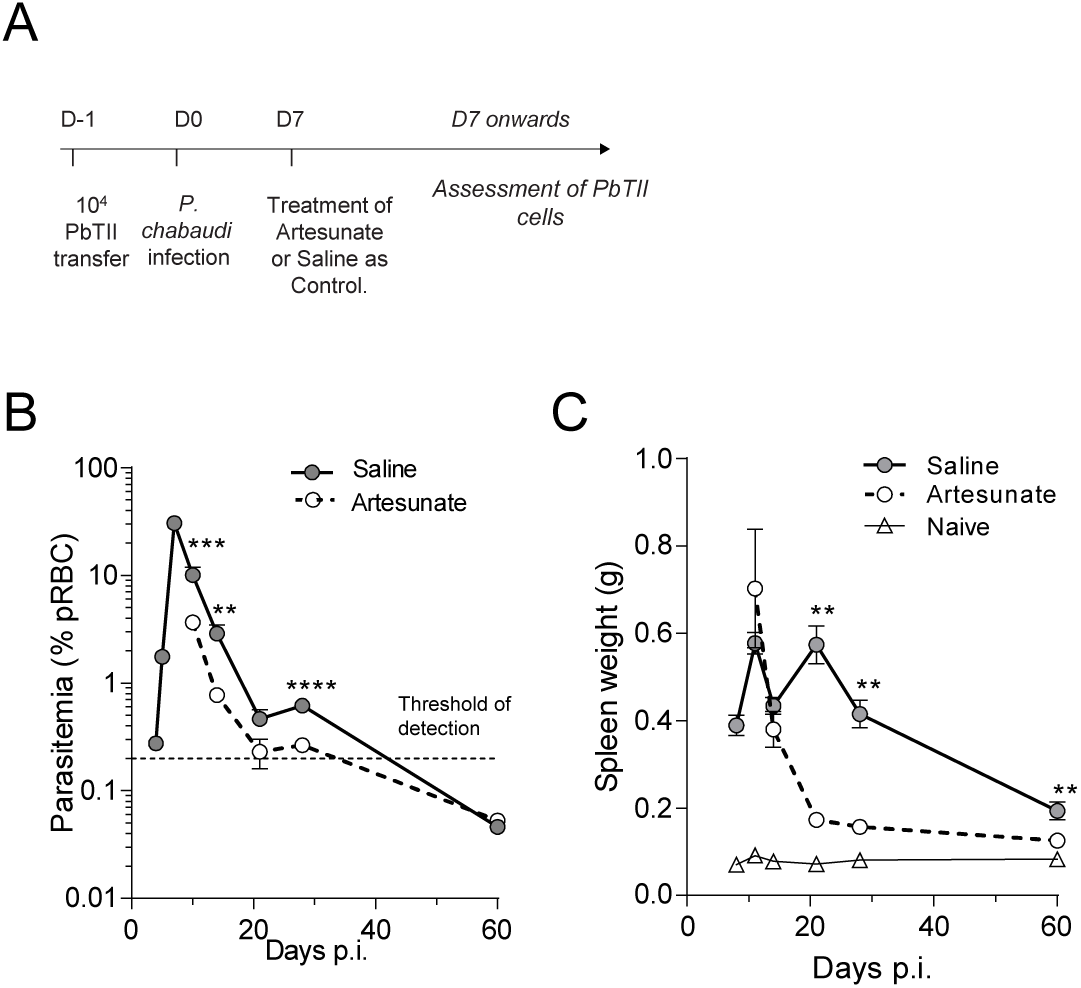
Model for studying CD4^+^ T cell memory during *in vivo Plasmodium* infection. **(A)** PbTII cells from a single donor were transferred to multiple recipients 24 hours prior to infection. Mice were then infected with *PcAS* pRBCs. At D7 *p.i.*, a group of mice were treated with artesunate, while another group of mice received saline treatment as vehicle control. Dosing regimen is described in Methods section. **(B)** Kinetics of parasitemia during *PcAS* infection in C57BL/6J mice. **(C)** Changes in spleen weight during *PcAS* infection. Data is either pooled from 4-5 independent experiments (B) or representative of 2-3 independent experiments (C) (n=5-6 mice/group/timepoint). Statistical analysis was performed between saline- and artesunate-treated groups for each individual timepoint using Mann-Whitney test. Data is presented as mean +/- SEM (B-C). *p< 0.05, **p< 0.01, ***p<0.001, ****p<0.0001

**Supp 2.**
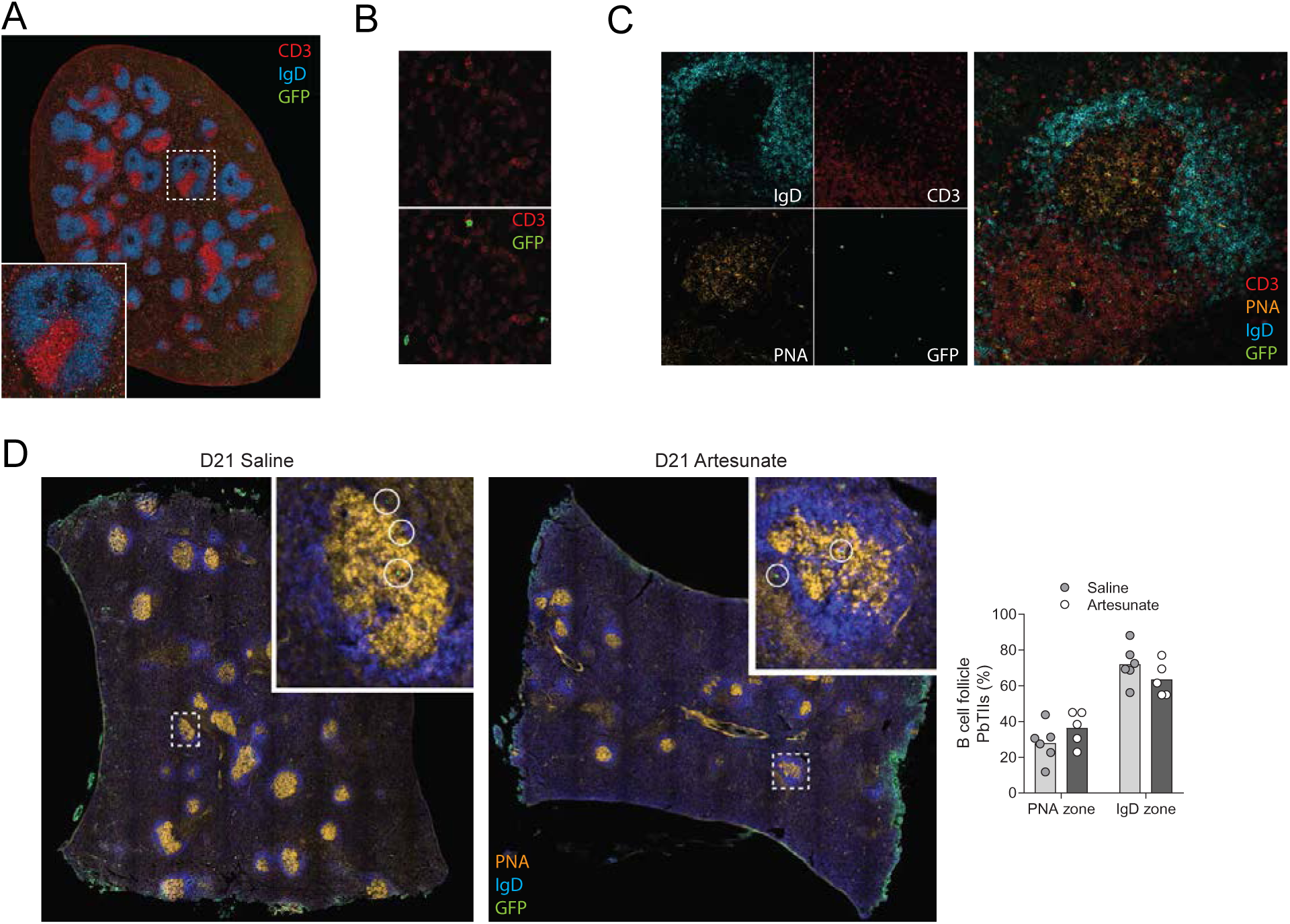
PbTII cells localise into T cell zones, B cell follicles and germinal centres (GC) during infection. **(A)** GFP^+^ PbTII cells were distributed throughout splenic B cell follicles and T cell zones at D7 *p.i*. (n=4). Graph showing proportion of PbTII cells localised within B cell (IgD) or T cell (CD3) zones. **(B)** Co-localisation of GFP and CD3 surface markers on CD4^+^ PbTII cells. **(C)** Standard microanatomical structures observed in saline-treated mice at D21 *p.i.*. **(D)** Localisation of PbTII cells within GCs and IgD+ naïve B cell follicle regions of saline or artesunate-treated mice at D21 *p.i.*. Graph showing proportion of PbTII cells localised within GCs or follicular regions of saline- (n=6) or artesunate- (n=5) treated mice. Images were acquired on an Aperio FL slide scanner (A, D) or a Zeiss 780-NLO point scanning confocal microscope (B-C). Statistical analysis was performed using two tailed t-test with Welch’s correction. Error bars illustrate 95% CI. **p < 0.01.

**Supp 3.**
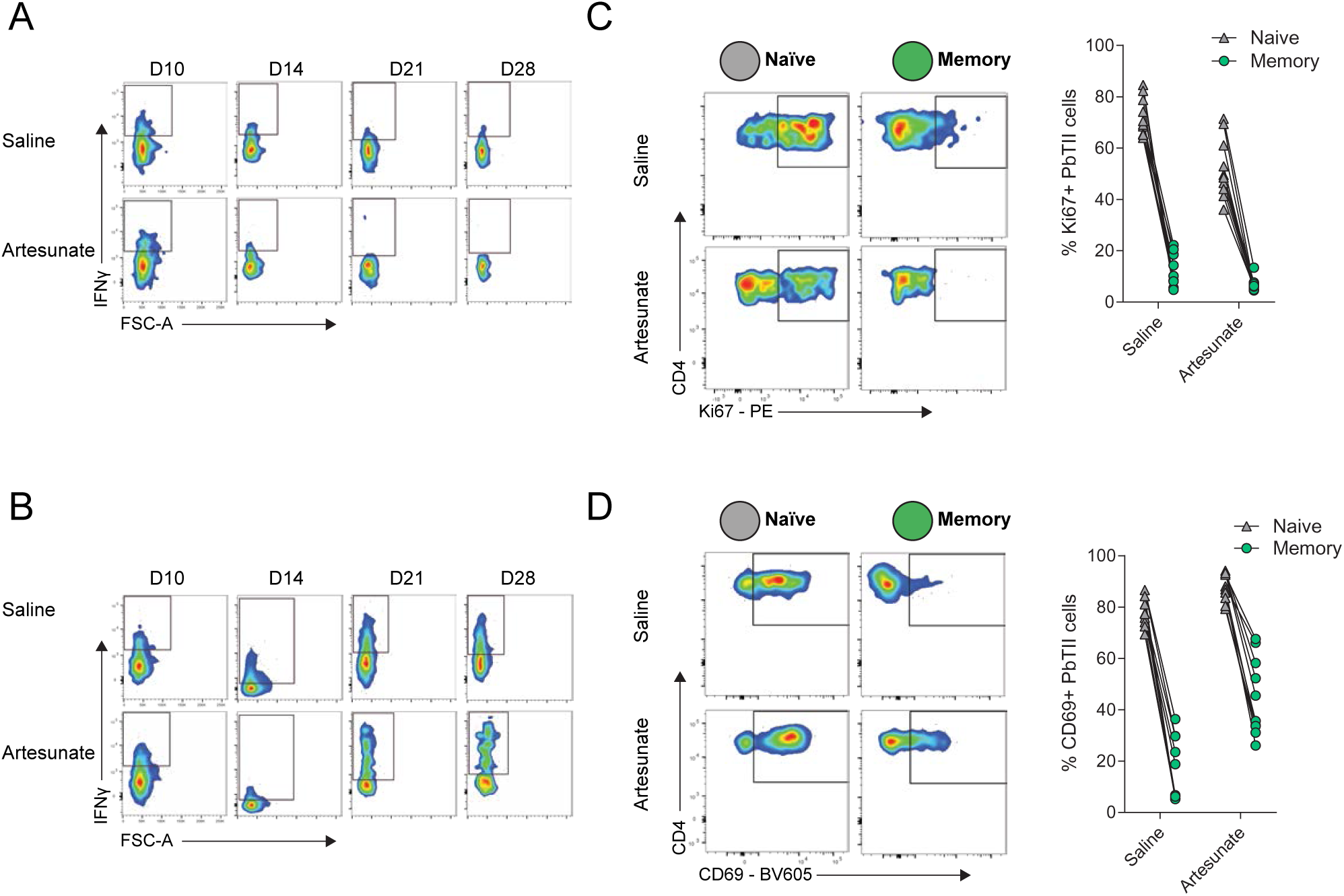
Memory PbTII cells are less proliferative compared to naïve cells during *in vivo* rechallenge. **(A-B)** Representative FACS plots for *ex vivo* IFNγ production assessed in Figure 2A-B, without any restimulation (A) or with restimulation with PMA/Ionomycin *in vitro* for 3 hours (B). **(C)** Representative FACS plots and graph showing proliferative marker Ki67 expression for memory (green) or naïve (grey) PbTII cells at 17 hours post-rechallenge. **(D)** Representative FACS plots and graph showing expression of early activation marker, CD69 for memory (green) or naïve (grey) PbTII cells at 17 hours post-rechallenge. Data is pooled from two experiments (n=5-6 mice/group/timepoint). Statistical analysis was performed using paired analysis. *p< 0.05, **p< 0.01, ***p<0.001, ****p<0.0001. Each dot represents an individual mouse.

**Supp 4.**
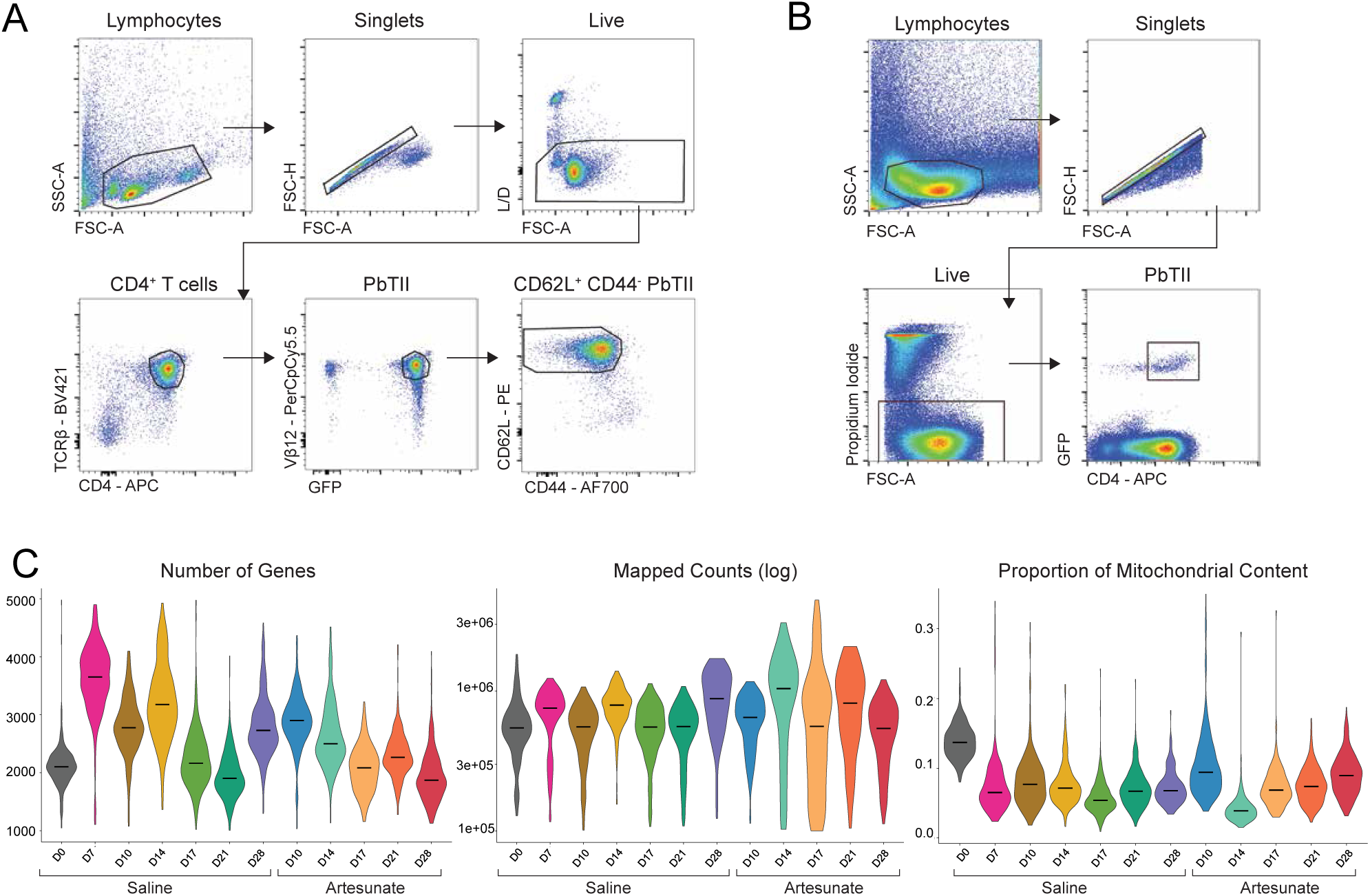
Quality checks for scRNA-seq assessment of memory PbTII cell differentiation. **(A)** FACS gating strategy for isolation of naïve (CD62L^+^ CD44^−^) donor PbTII cells for transfer into recipients, 24 hours prior to infection. A subset of these cells was also used for scRNA-seq assessment for PbTII responses at D0 *p.i.*. Cells were then either sorted as single-cells onto 384-well plates for Smart-seq2 analysis (Figure 3-7) or sorted as single-cells using the 10x Chromium platform (Figure S6 and Figure S16). **(B)** FACS gating strategy for isolation of PbTII cells from D7 *p.i.* onwards for scRNA-seq assessment for either Smart-seq2 (Figure 3-7) or 10x Chromium assessment (Figure S6 and Figure S16). **(C)** Distribution of PbTII cells after filtering for number of genes (1000<nGene< 5000), mapped counts (> 100,000) and proportion of mitochondrial content (<0.35).

**Supp 5.**
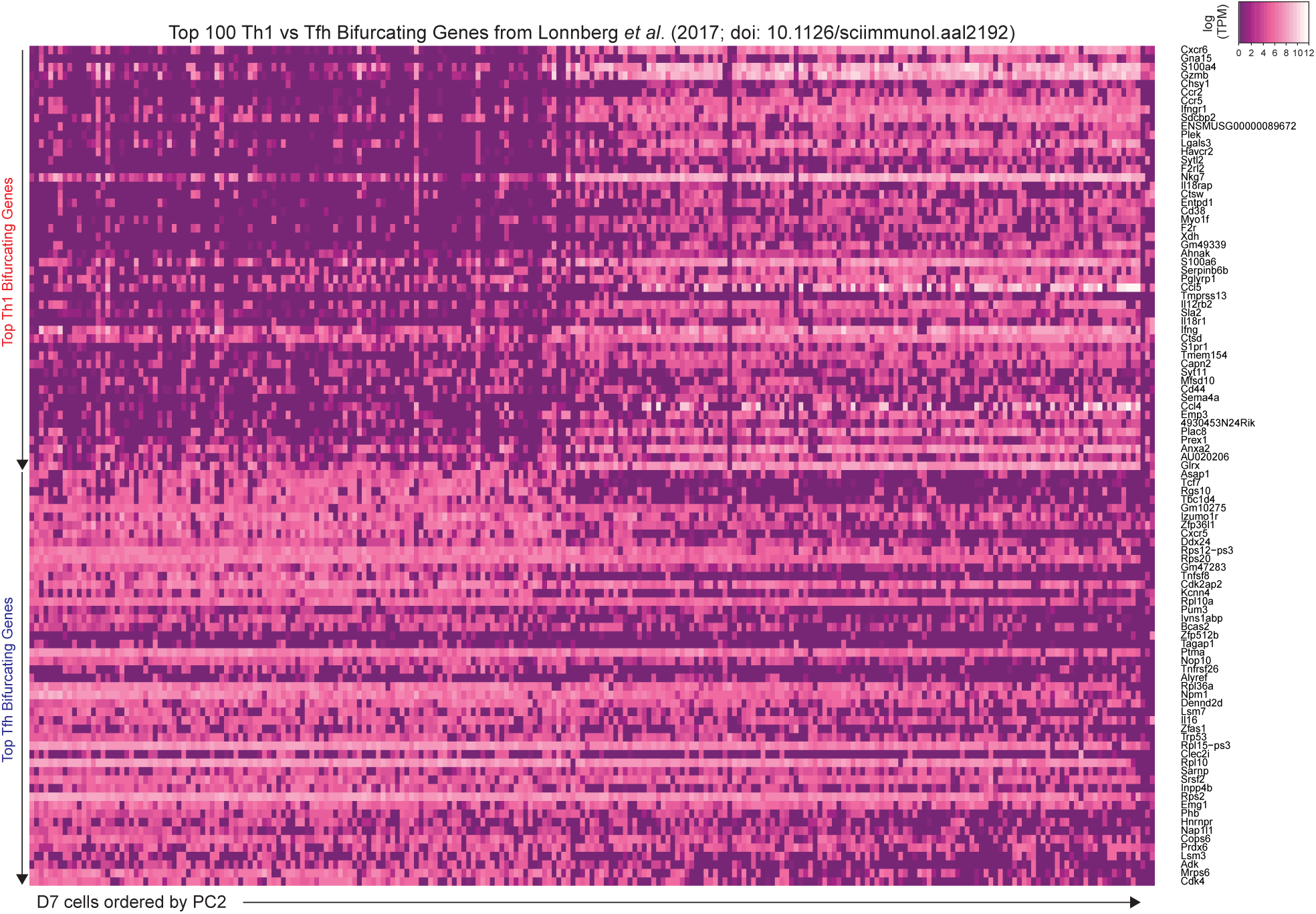
Clusters of PbTII cells at D7 p.i. corresponded to either Th1 or Tfh effector fate. Heatmap describing expression of top 50 Th1 or Tfh-bifurcating genes derived from Lönnberg et al. (2017) (Lonnberg et al., 2017) for PbTII cells from D7 *p.i.*. Genes are ranked in descending order according to their bifurcation statistics (y-axis), while cells are ordered along their position based on PC2 coordinates (Figure 3B).

**Supp 6.**
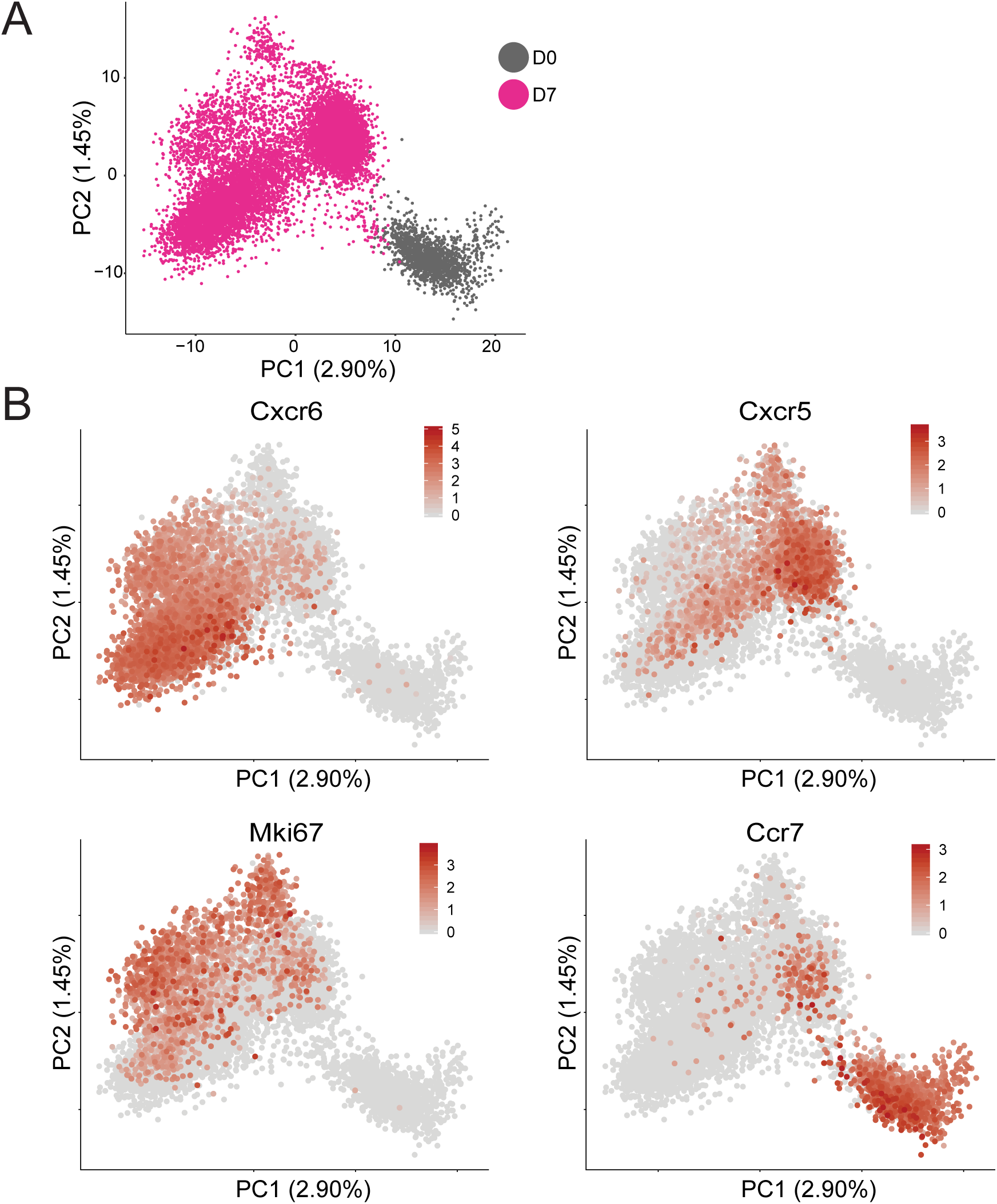
Effector PbTII cells at D7 p.i. largely corresponded to Th1 or Tfh effector fate with a minor population of highly-proliferative cells. **(A)** Experiment was designed as in Figure 3A but performed on the 10x Chromium platform. Naïve (CD62L^+^ CD44^−^) donor PbTII cells from one donor were transferred into multiple recipients, 24 hours prior to infection. A subset of these cells was also used for scRNA-seq assessment for PbTII responses at D0 *p.i.* Five mice were pooled for isolating splenic PbTII cells at D7 *p.i.* and sorted as single cells using the 10x Chromium platform. PCA of PbTII cells from D0 (1496 cells) and D7 p.i. (10788 cells) was performed using 2207 highly variable genes as input after regressing out number of unique molecular identifiers (nUMI). **(B)** Visualisation of *Cxcr6, Cxcr5, Mki67* and *Ccr7* expression on PbTII cells as in (A).

**Supp 7.**
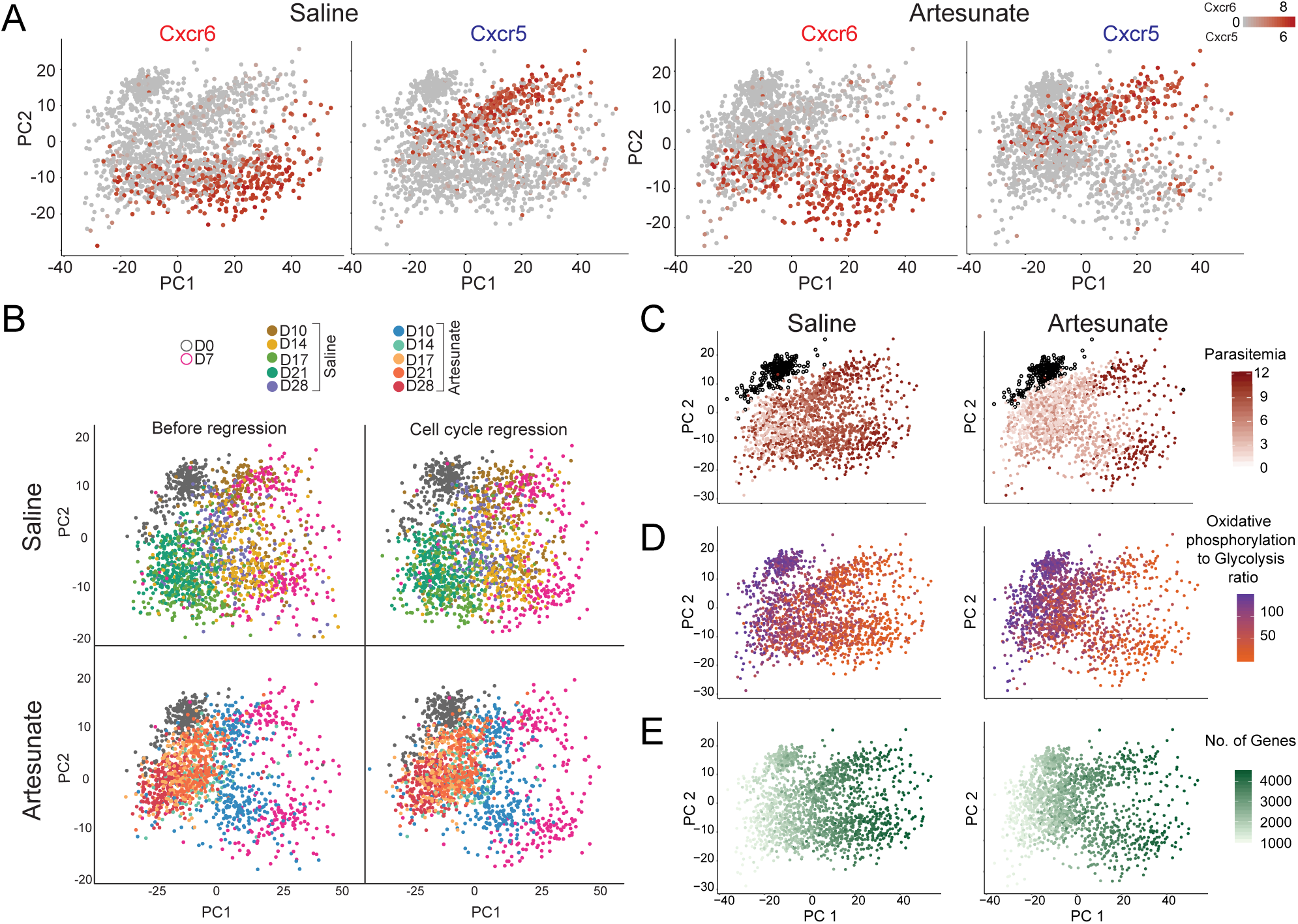
Characterisation of transcriptomes of PbTII cells during effector to memory transition. **(A)** PCA (as described in Figure 3C) plots coloured with *Cxcr6* and *Cxcr5* expression. **(B)** PCA showed minimal effect of cell cycle regression. Cell cycle regression was performed based on 226 cell cycle genes derived from Cyclebase (Santos et al., 2015), using Seurat’s built-in regressing out function. **(C)** PCA plots coloured with parasitemia of the specific donor mouse used for isolating PbTII cells at each timepoint/group. **(D)** PCA plots coloured with the ratio of oxidative phosphorylation levels to glycolysis levels, expressed per cell. Oxidative phosphorylation levels was the summative expression of 28 genes (GO: 0006119) while glycolysis levels was the summative expression of 31 genes (GO: 0006096). **(E)** PCA plots coloured with the total number of genes expressed per cell.

**Supp 8.**
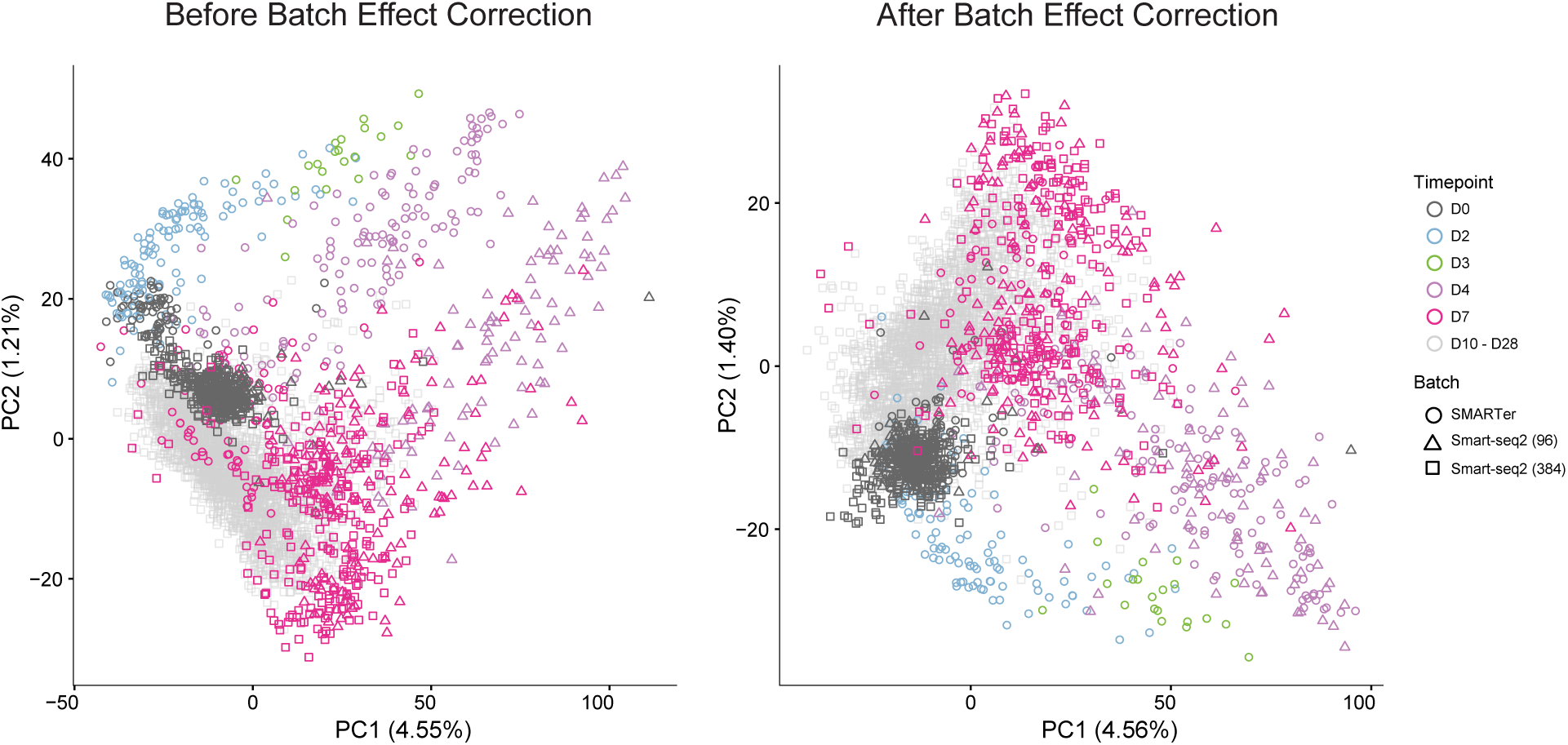
Batch correction of the entire PbTII time series revealed progression of cellular responses from naivety to memory. The current PbTII dataset of D0, D7, D10*, D14*, D17*, D21* and D28* *p.i.* cells were combined with our previous datasets (Lonnberg et al., 2017) containing PbTII cells isolated from D0 to D7 *p.i.* (SMARTer batch: D0, D2, D3, D4, D7 *p.i.*; Smart-seq2 (96) batch: D0, D4, D7 *p.i.*). PCA showing the entire time series, with shapes denoting the different experiment batches, before (left) and after (right) batch effect correction as described in methods. (*) = samples were isolated either from saline or artesunate-treated mice.

**Supp 9.**
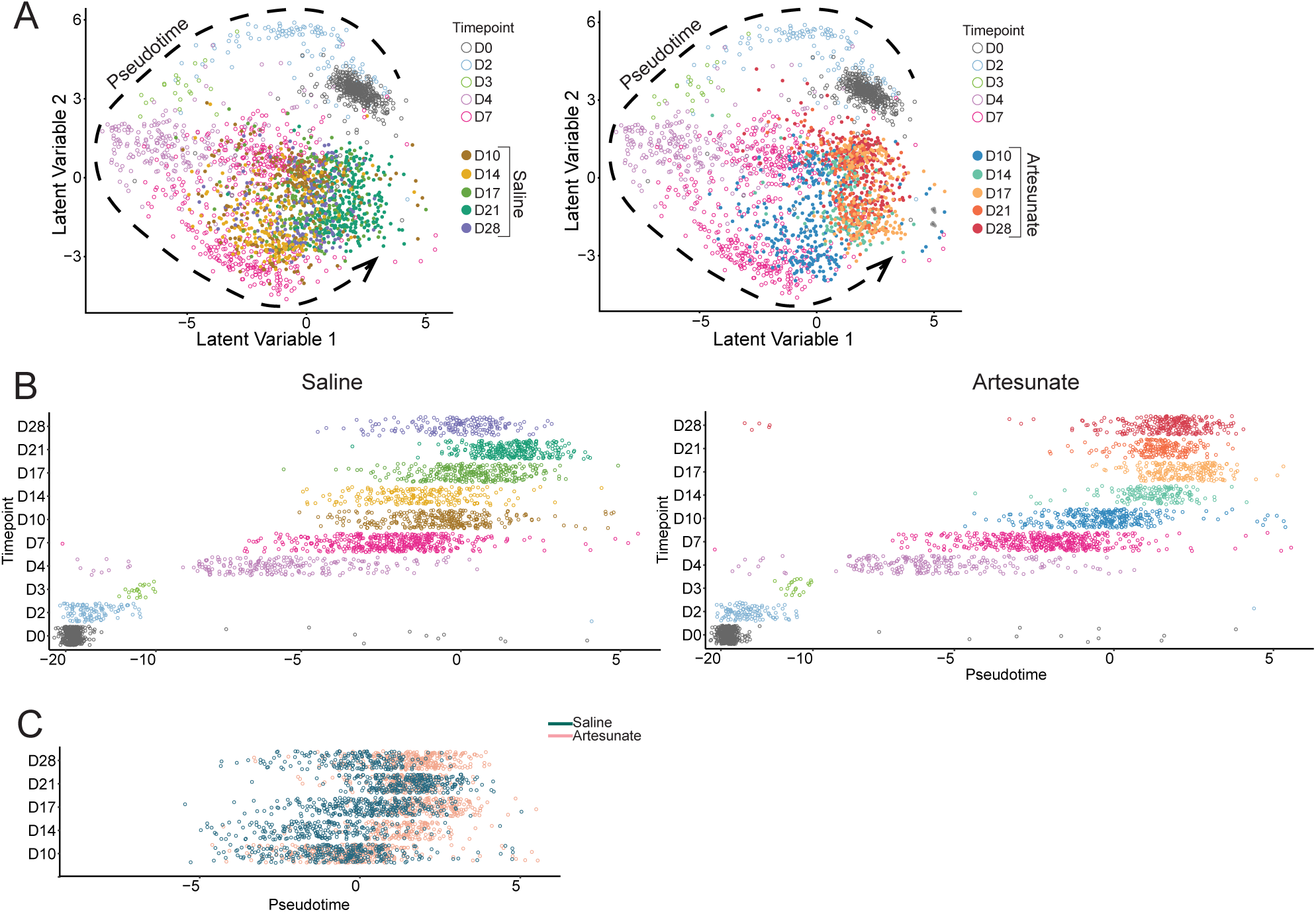
Temporal ordering of PbTII cells from naivety to memory. **(A)** BGPLVM analysis was performed on the full time series from the combined PbTII datasets (Figure S8), and 2D representation was plotted based on treatment groups (left=saline; right=artesunate). D0, D2, D3, D4 and D7 p.i. cells were replicated on each sub-plot. Dashed lines denote pseudotime inference as described in methods. **(B)** Scatter plots of PbTII cells from each timepoint along pseudotime (left=saline; right=artesunate). **(C)** Comparison of PbTII cells from each timepoint along pseudotime, for saline and artesunate-treated groups.

**Supp 10.**
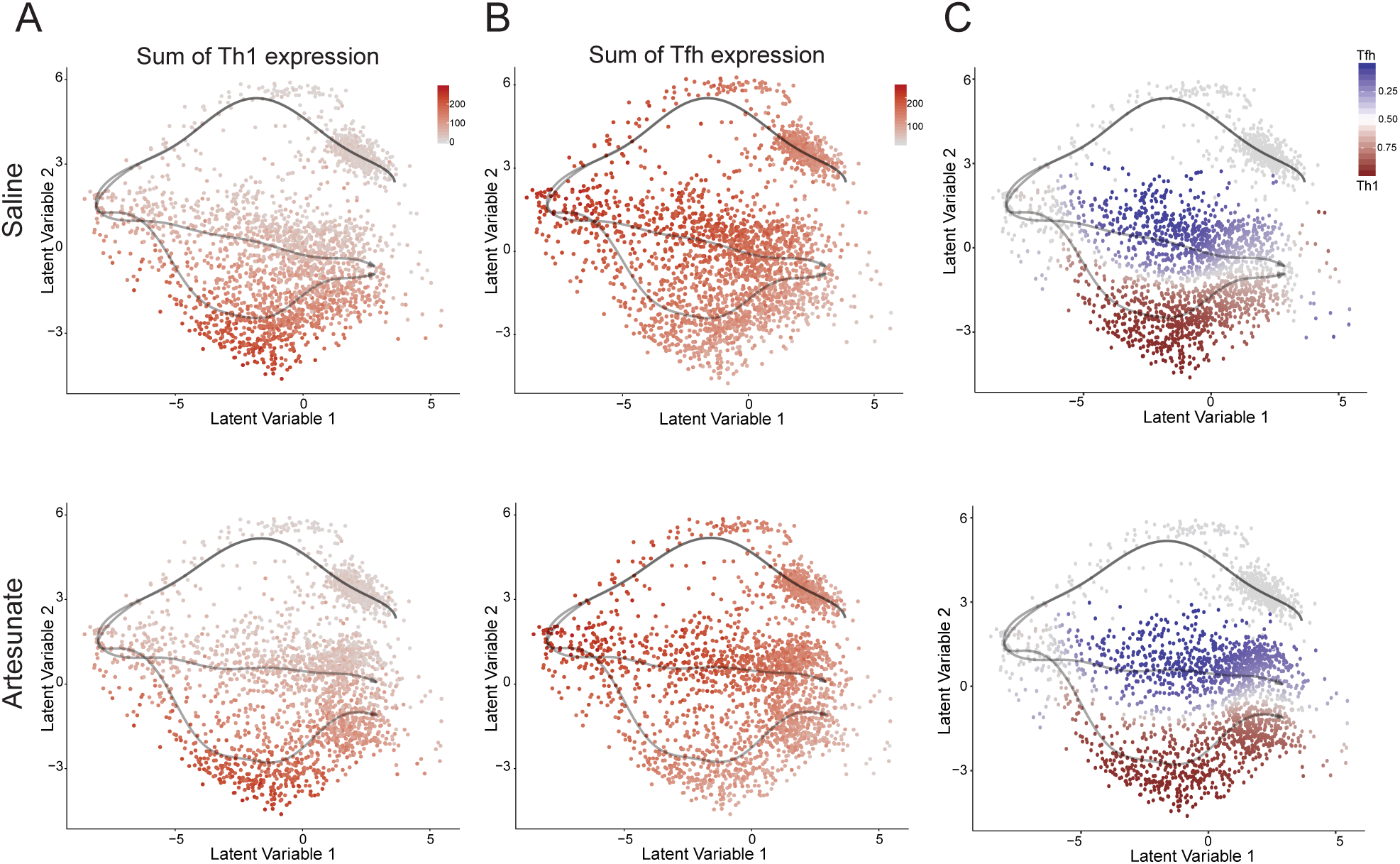
PbTII Th1/Tfh effector fates are retained during progression into memory PbTII cells. **(A)** Visualisation of Th1 signature (summative expression of top 50 Th1-bifurcating genes derived from Lönnberg *et al*. (2017) (Lonnberg et al., 2017) on BGPLVM 2D representation. **(B)** Visualisation of Tfh signature (summative expression of top 50 Th1-bifurcating genes derived from Lönnberg *et al*. (2017) (Lonnberg et al., 2017) on BGPLVM 2D representation. **(C)** Visualisation of Th1/Tfh assignment probability as calculated by OMGP analysis using GPfates on BGPLVM 2D representation.

**Supp 11.**
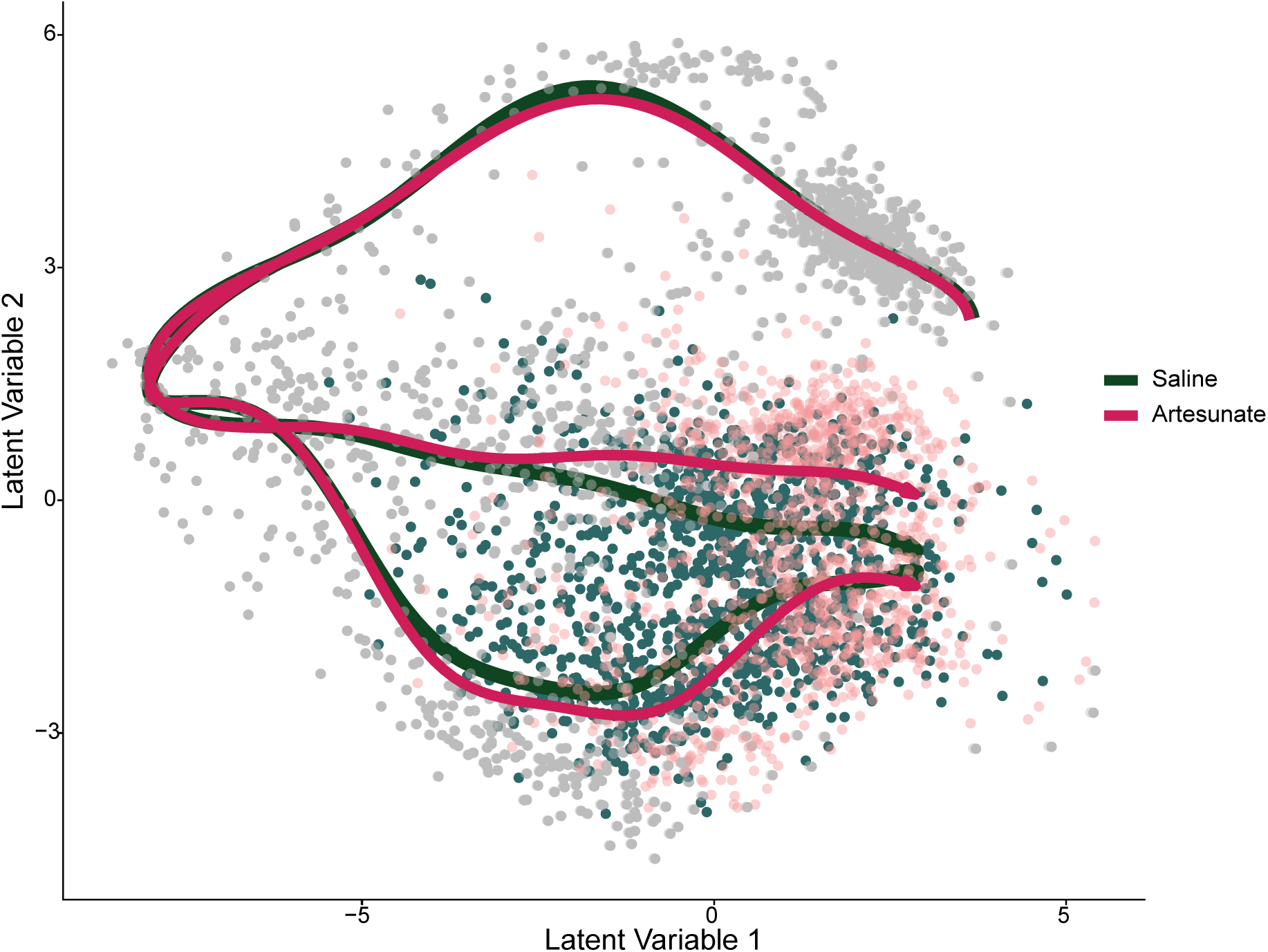
PbTII cells from late timepoints from the artesunate-treated group are positioned later along pseudotime compared to saline-treated cells. Positional comparison of saline versus artesunate-treated PbTII cells, from D10 *p.i.* onwards, on BGPLVM 2D representation. Separate OMGP trends are shown for saline and artesunate groups.

**Supp 12.**
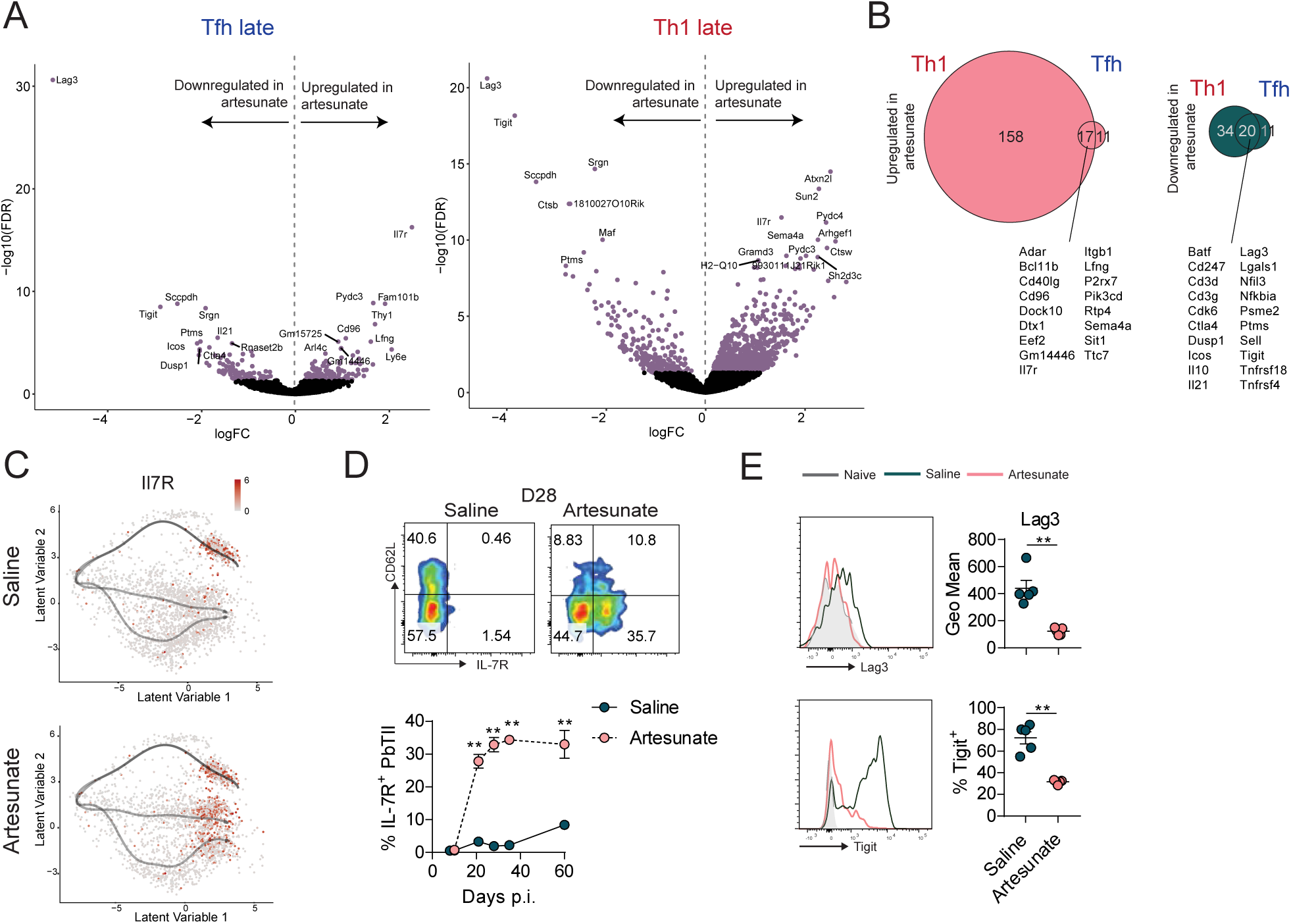
Memory PbTII cells developed in the absence of persisting infection predicted better functional memory responses. **(A)** Identification of differentially-expressed genes from late cells (positioned at pseudotime>0.9; Figure 4E) between artesunate- and saline-treated mice for Tfh branch and Th1 branch (Figure S10C), respectively. Genes with false discovery rate (FDR) less than 0.05 are coloured with lavender. **(B)** Venn diagrams showing the number of differentially expressed genes involved in immune system process (GO:0002376). Overlapping list of genes between (left) upregulated in artesunate-treated late cells on Th1 branch and upregulated in artesunate-treated late cells on Tfh branch, and (right) upregulated in saline-treated late cells on Th1 branch and upregulated in saline-treated late cells on Tfh branch. **(C)** Visualisation of *Il7r* expression on BGPLVM 2D representation. **(D)** Protein level measurement of surface IL7R on PbTII cells. Representative FACS plots and graph showing the kinetics of IL7R expression over time. Data is representative of 2 independent experiments. **(E)** Representative FACS plots and graph showing surface expression of LAG3 and TIGIT on PbTII cells at D28 *p.i.*. Experiment was performed once. Data is presented as mean +/- SEM (n=5-6 mice/group/experiment). Statistical analysis was performed using Mann-Whitney test. **p< 0.01

**Supp 13.**
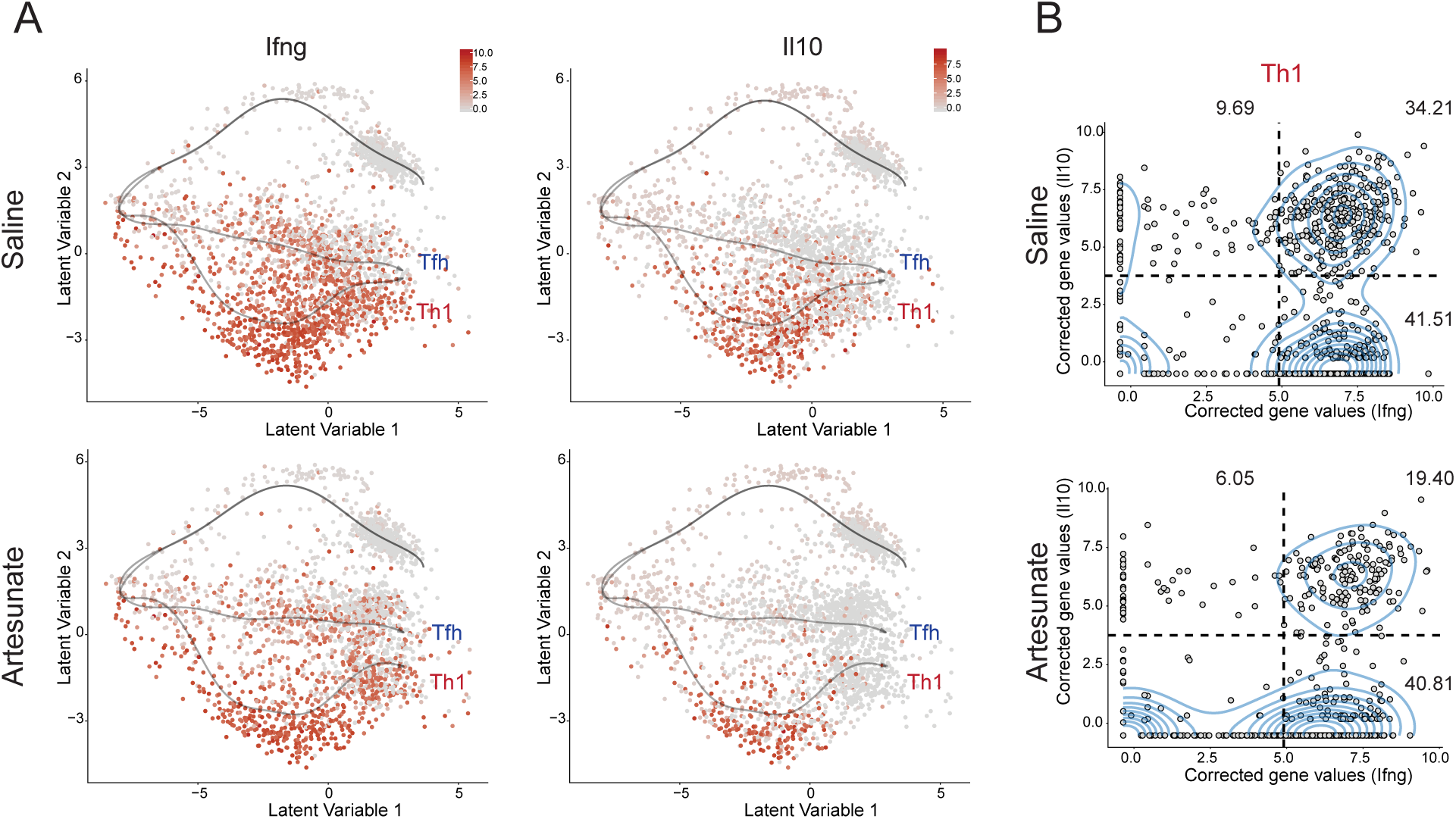
A subset of Th1 PbTII cells developed into Tr1 cells. **(A)** Visualisation of *Il10* and *Ifng* on 2D BGPLVM representation. OMGP trajectories representing Th1 or Tfh branch are shown. **(B)** Co-expression of *Il10* and *Ifng* along pseudotime for all cells annotated for Th1 (Figure S10C). Tr1 cells are annotated as those cells expressing *Il10* and *Ifnγ* above the threshold drawn (cells expressing both *Il10*≥4.9 and *Ifng*≥3.75 corrected gene values), according to kernel density estimation.

**Supp 14.**
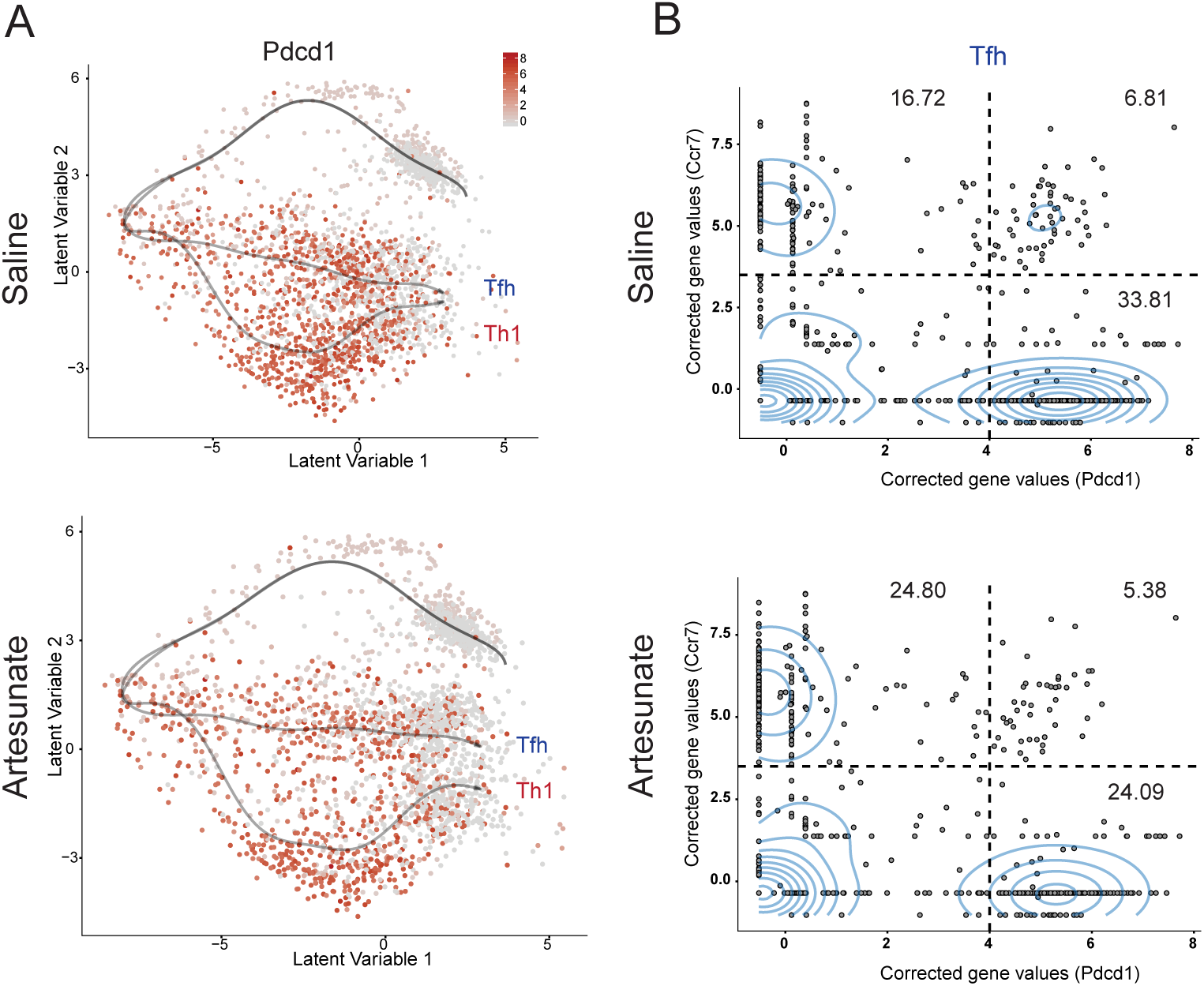
Characterisation of GC Tfh cells in memory PbTII cells. **(A)** Visualisation of *Pdcd1* on 2D BGPLVM representation. OMGP trajectories representing Th1 or Tfh branch are shown. **(B)** Co-expression of *Pdcd1* and *Ccr7* along pseudotime, for all cells annotated for the Tfh branch according to (Figure S10C). Kernel density estimation failed to estimate a dense population of *Pdcd1^+^ Ccr7^+^* cells, hence a threshold was estimated where cells express both *Pdcd1* ≥ 4.0 and *Ccr7* ≥ 3.50 corrected gene values.

**Supp 15.**
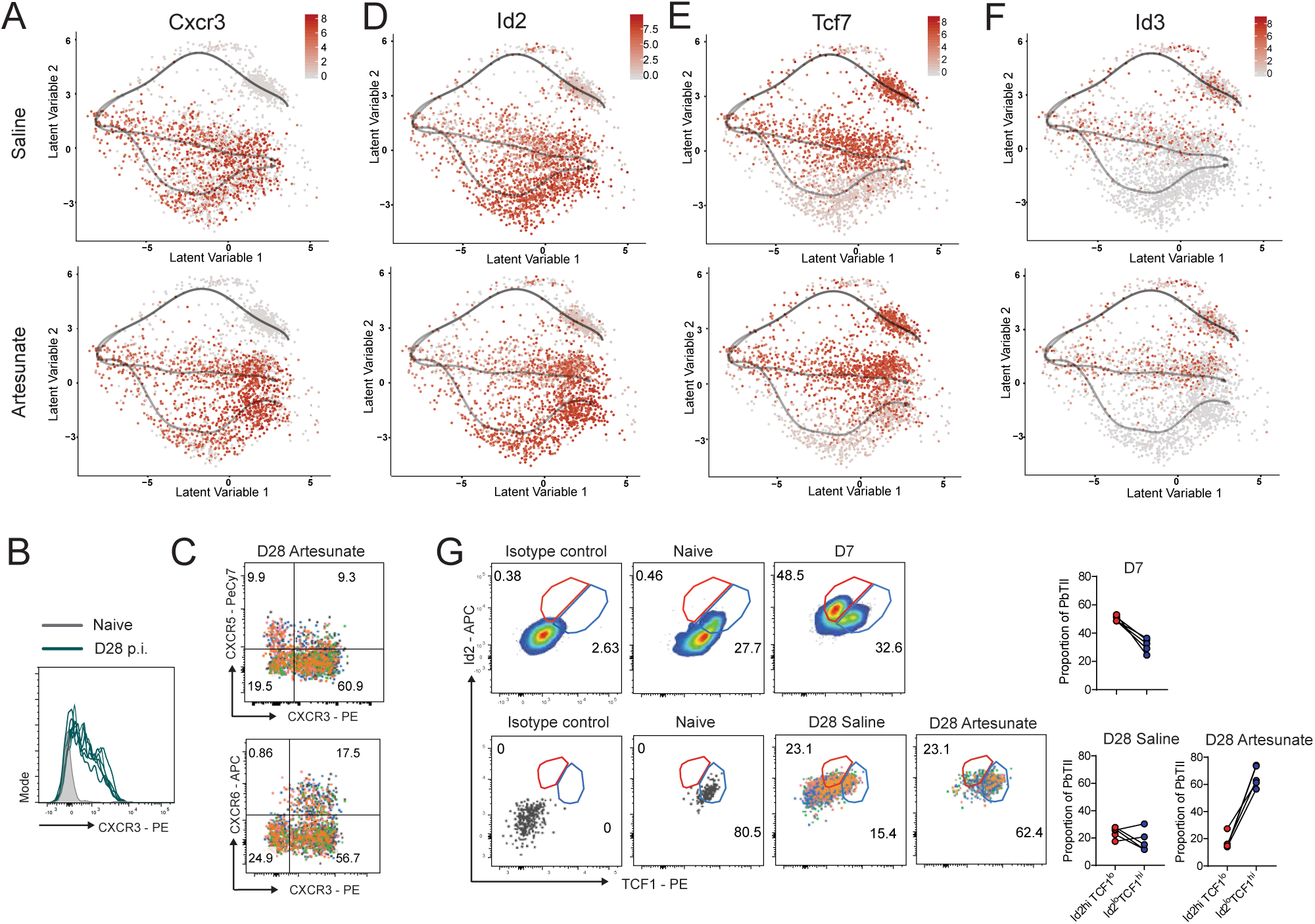
Expression of memory-associated genes on PbTII cells. **(A)** Visualisation of *Cxcr3* on 2D BGPLVM representation. **(B)** FACS plot showing surface CXCR3 expression on PbTII cells at naïve versus saline-treated mice at D28 p.i. Each line represents an individual mouse. Data is representative of 3 independent experiments (n=5-6 mice). **(C)** FACS plot showing surface CXCR3 versus CXCR5 or CXCR6 expression on PbTII cells for artesunate-treated mice at D28 p.i. Each colour represents an individual mouse. Experiment is performed once. **(D)** Visualisation of *Id2* on 2D BGPLVM representation. **(E)** Visualisation of *Tcf7* on 2D BGPLVM representation. **(F)** Visualisation of *Id3* on 2D BGPLVM representation. **(G)** Representative FACS plot and graph showing protein expression of ID2 and TCF1 on PbTII cells at D7 *p.i.* (top). FACS plots and graph showing protein expression of ID2 and TCF1 on PbTII cells from saline- or artesunate-treated mice at D28 *p.i.* (bottom, each colour on the FACS plot represent an individual mouse).

**Supp 16.**
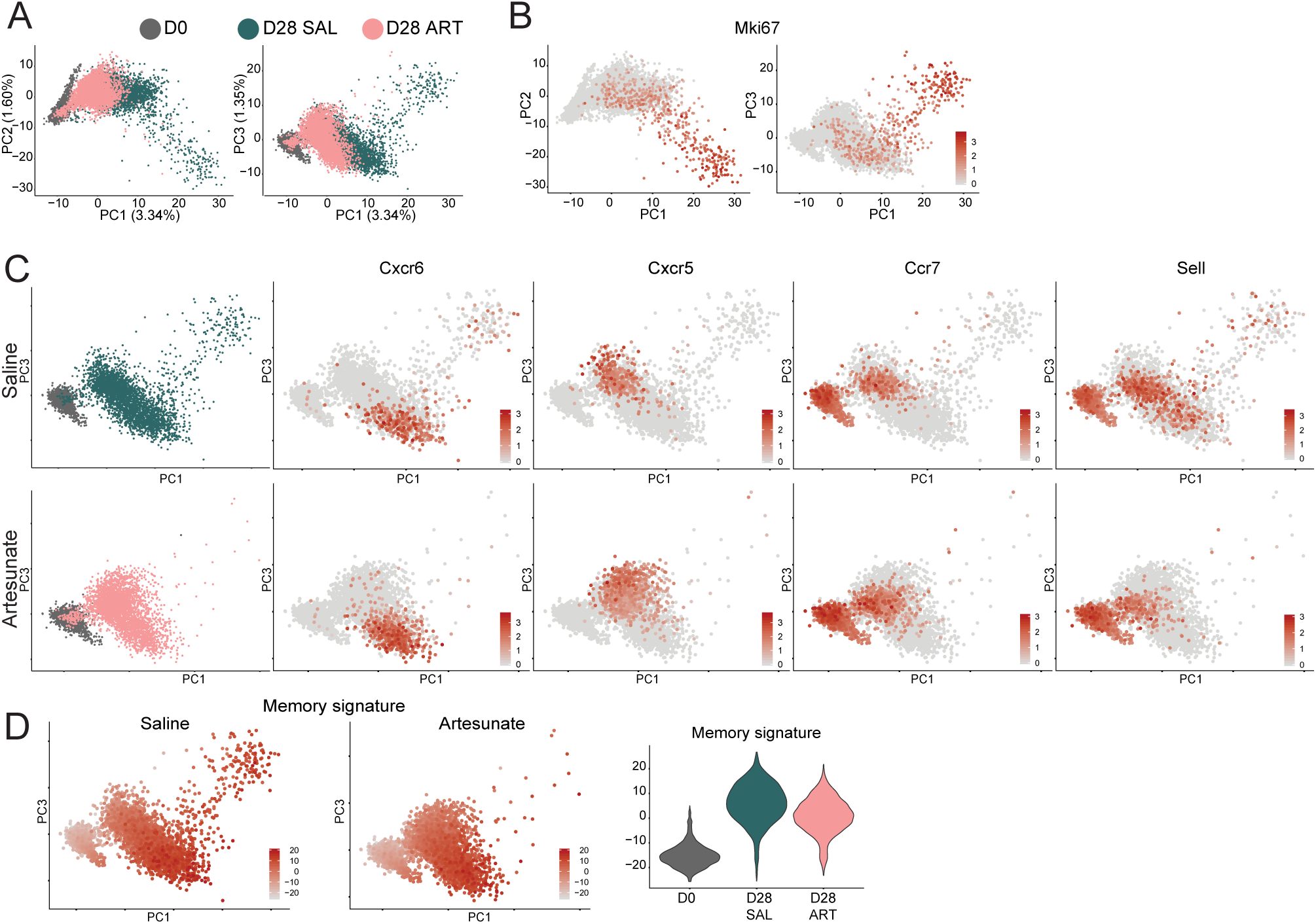
Expression of effector-associated and memory-associated genes on memory PbTII cells. **(A)** Experiment was designed as in Figure 3A but performed on the 10x Chromium platform. 15 mice per treatment group were pooled for isolating splenic PbTII cells at D28 *p.i.* and sorted as single-cells using the 10xChromium platform. PCA analysis of D0 (1496 cells), D28 SAL (saline-treated cells) (3300 cells) and D28 ART (artesunate-treated cells) (3472 cells). PCA was performed based on 1515 highly variable genes, after regression out nUMI. **(B)** Visualisation of *Mki67* expression on PCA plot. **(C)** Visualisations of *Cxcr6, Cxcr5, Sell, Ccr7* expression on PCA plots. **(D)** Visualisation of memory-associated signature (summative expression of top 20 pseudotime correlated genes from artesunate-treated group described in Figure 7E). Comparison of expression for memory-associated signature (Figure S16B) between D0, D28 SAL and D28 ART.

**Supp 17.**
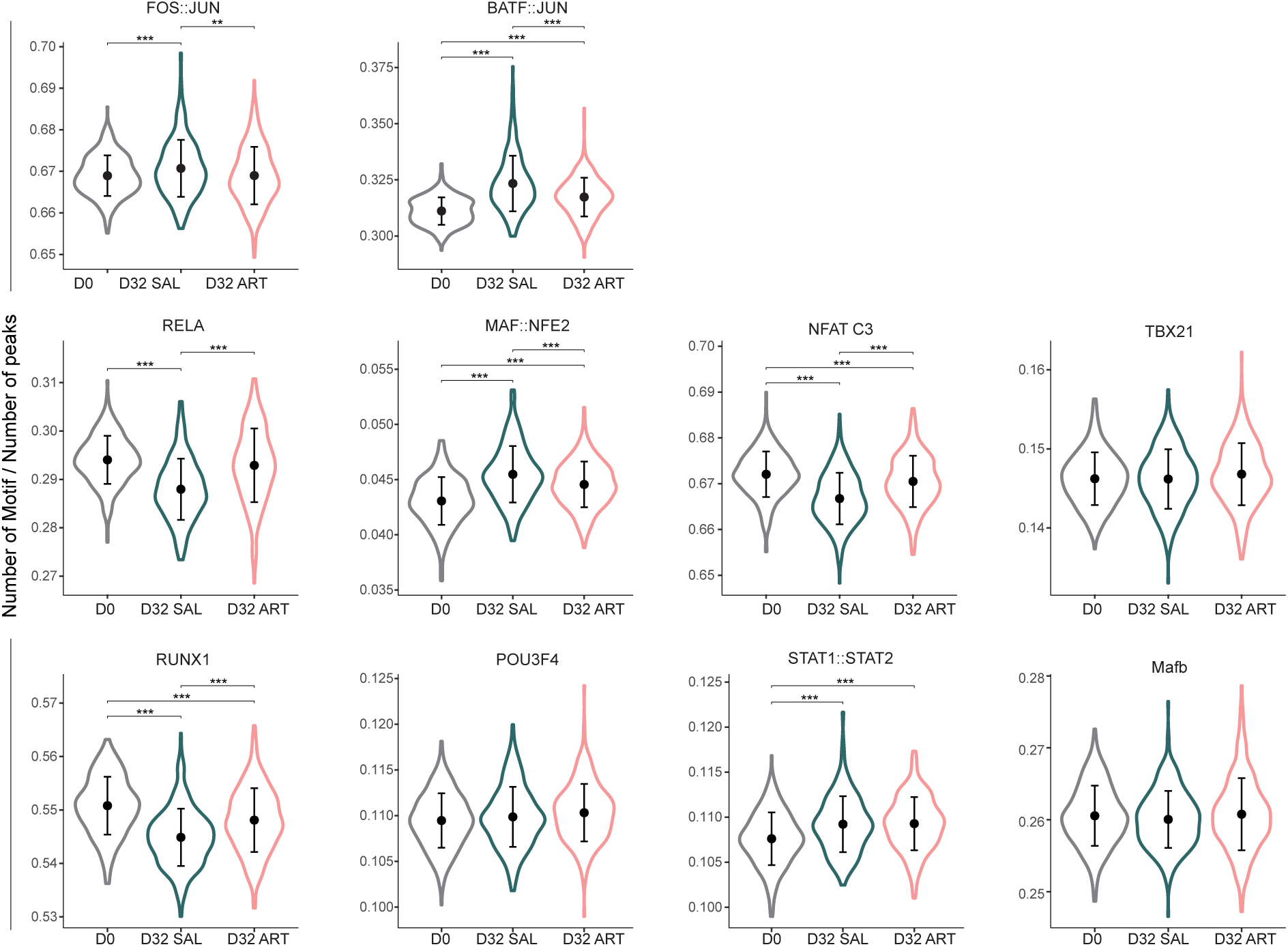
Partial resetting of chromatin landscape accompanied memory transition. Changes in chromatin accessibility for naïve (D0) PbTII cells, saline and artesunate-treated cells at D32 *p.i.* associated with different transcription factors.

