## Supplementary Table 3 for "Transcriptome Dynamics Reveals Progressive Transition from Effector to Memory in CD4^+^ T cells"

| Table S3: **Top 50 variable motifs between D0, D32 saline and D32 artesunate samples** | JASPAR Matrix ID | Variability | p_value | p_value_adj | Family | Motif |
| --- | --- | --- | --- | --- | --- | --- |
| 1 | MA0491.1_JUND | 3.601197198 | 0 | 0 | Jun-related factors | 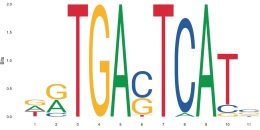 |
| 2 | MA0489.1_JUN(var.2) | 3.562425843 | 0 | 0 | Jun-related factors | 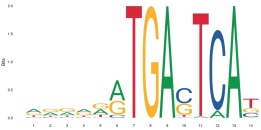 |
| 3 | MA0476.1_FOS | 3.545758842 | 0 | 0 | Fos-related factors | 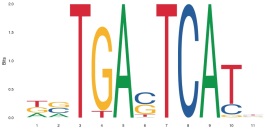 |
| 4 | MA0099.2_FOS::JUN | 3.532062621 | 0 | 0 | Fos-related factors::Jun-related factors | 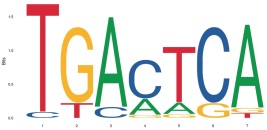 |
| 5 | MA0490.1_JUNB | 3.530172127 | 0 | 0 | Jun-related factors | 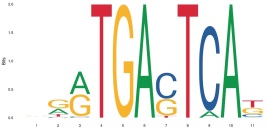 |
| 6 | MA0477.1_FOSL1 | 3.499961764 | 0 | 0 | Fos-related factors | 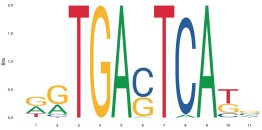 |
| 7 | MA0462.1_BATF::JUN | 3.454645932 | 0 | 0 | B-ATF-related factors::Jun-related factors | 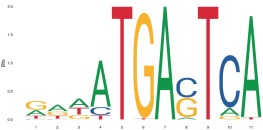 |
| 8 | MA0478.1_FOSL2 | 3.399126553 | 0 | 0 | Fos-related factors | 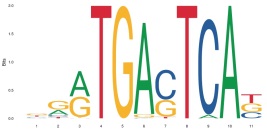 |
| 9 | MA0655.1_JDP2 | 3.294605143 | 0 | 0 | Fos-related factors | 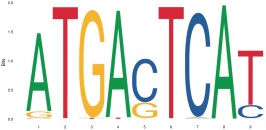 |
| 10 | MA0841.1_NFE2 | 2.871549669 | 0 | 0 | Jun-related factors | 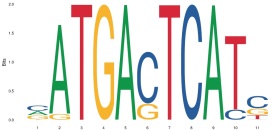 |
| 11 | MA0107.1_RELA | 2.115365487 | 0 | 0 | NF-kappaB-related factors | 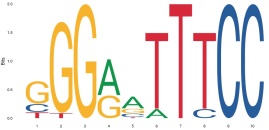 |
| 12 | MA0101.1_REL | 2.063878904 | 0 | 0 | NF-kappaB-related factors | 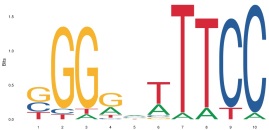 |
| 13 | MA0523.1_TCF7L2 | 1.927345648 | 2.20E-304 | 8.59E-303 | TCF-7-related factors | 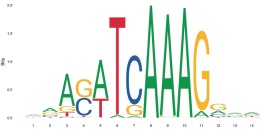 |
| 14 | MA0501.1_MAF::NFE2 | 1.894701223 | 6.93E-285 | 2.51E-283 | Maf-related factors::Jun-related factors | 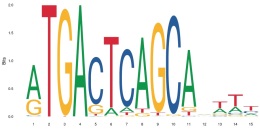 |
| 15 | MA0778.1_NFKB2 | 1.863439416 | 9.22E-267 | 3.12E-265 | NF-kappaB-related factors | 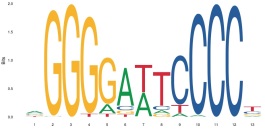 |
| 16 | MA0139.1_CTCF | 1.853756801 | 2.94E-261 | 9.32E-260 | More than 3 adjacent zinc finger factors | 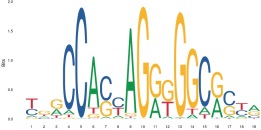 |
| 17 | MA0150.2_Nfe2l2 | 1.809297307 | 1.18E-236 | 3.52E-235 | Jun-related factors | 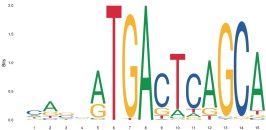 |
| 18 | MA0768.1_LEF1 | 1.73141025 | 3.12E-196 | 8.78E-195 | TCF-7-related factors | 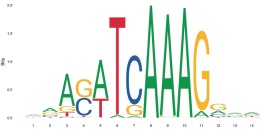 |
| 19 | MA0591.1_Bach1::Mafk | 1.691382648 | 8.34E-177 | 2.22E-175 | Jun-related factors::Maf-related factors | 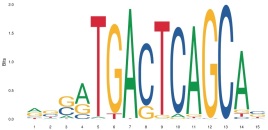 |
| 20 | MA0652.1_IRF8 | 1.689248499 | 8.53E-176 | 2.16E-174 | Interferon-regulatory factors | 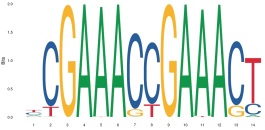 |
| 21 | MA0653.1_IRF9 | 1.672509503 | 5.75E-168 | 1.39E-166 | Interferon-regulatory factors | 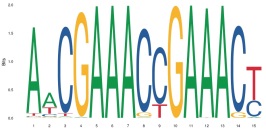 |
| 22 | MA0769.1_Tcf7 | 1.651409826 | 2.49E-158 | 5.73E-157 | TCF-7-related factors | 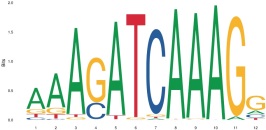 |
| 23 | MA0625.1_NFATC3 | 1.647280228 | 1.78E-156 | 3.93E-155 | NFAT-related factors | 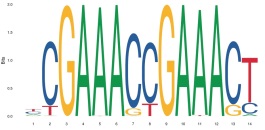 |
| 24 | MA0495.1_MAFF | 1.61648941 | 5.86E-143 | 1.24E-141 | Maf-related factors | 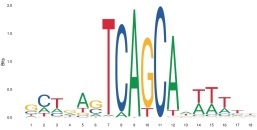 |
| 25 | MA0105.4_NFKB1 | 1.577694735 | 9.94E-127 | 2.02E-125 | NF-kappaB-related factors | 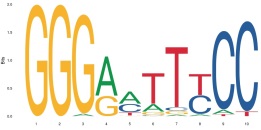 |
| 26 | MA0496.1_MAFK | 1.547446217 | 1.06E-114 | 2.06E-113 | Maf-related factors | 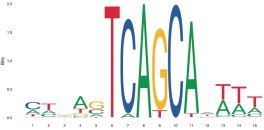 |
| 27 | MA0688.1_TBX2 | 1.531902586 | 9.74E-109 | 1.83E-107 | TBX2-related factors | 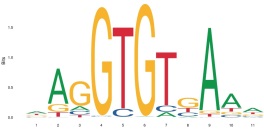 |
| 28 | MA0800.1_EOMES | 1.488777924 | 5.75E-93 | 1.04E-91 | TBrain-related factors | 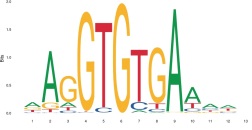 |
| 29 | MA0690.1_TBX21 | 1.481429285 | 2.15E-90 | 3.75E-89 | TBrain-related factors | 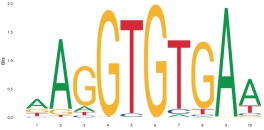 |
| 30 | MA0772.1_IRF7 | 1.462974218 | 4.40E-84 | 7.44E-83 | Interferon-regulatory factors | 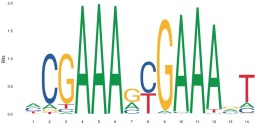 |
| 31 | MA0805.1_TBX1 | 1.456304835 | 7.45E-82 | 1.22E-80 | TBX1-related factors | 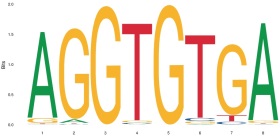 |
| 32 | MA0051.1_IRF2 | 1.443990526 | 8.19E-78 | 1.30E-76 | Interferon-regulatory factors | 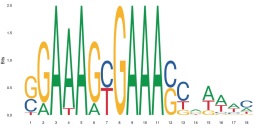 |
| 33 | MA0802.1_TBR1 | 1.433071527 | 2.60E-74 | 4.00E-73 | TBrain-related factors | 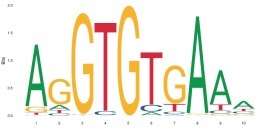 |
| 34 | MA0002.2_RUNX1 | 1.428996109 | 5.05E-73 | 7.54E-72 | Runt-related factors | 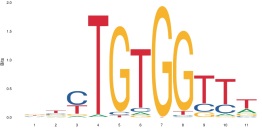 |
| 35 | MA0801.1_MGA | 1.427447634 | 1.55E-72 | 2.18E-71 | TBX6-related factors | 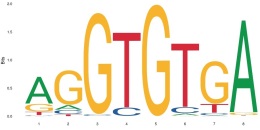 |
| 36 | MA0803.1_TBX15 | 1.427447634 | 1.55E-72 | 2.18E-71 | TBX1-related factors | 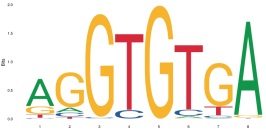 |
| 37 | MA0789.1_POU3F4 | 1.411995191 | 9.15E-68 | 1.25E-66 | POU domain factors | 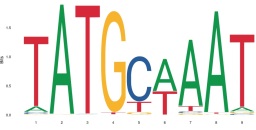 |
| 38 | MA0517.1_STAT1::STAT2 | 1.411856687 | 1.01E-67 | 1.35E-66 | STAT factors::STAT factors | 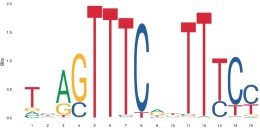 |
| 39 | MA0785.1_POU2F1 | 1.411579002 | 1.22E-67 | 1.59E-66 | POU domain factors | 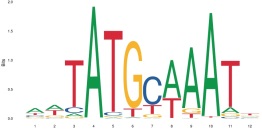 |
| 40 | MA0645.1_ETV6 | 1.402752113 | 5.52E-65 | 7.00E-64 | Ets-related factors | 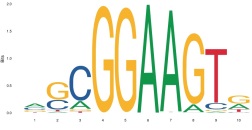 |
| 41 | MA0792.1_POU5F1B | 1.400256979 | 3.04E-64 | 3.76E-63 | POU domain factors | 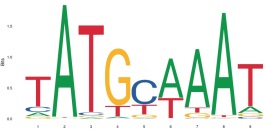 |
| 42 | MA0117.2_Mafb | 1.392210731 | 7.02E-62 | 8.47E-61 | Maf-related factors | 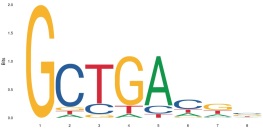 |
| 43 | MA0689.1_TBX20 | 1.388599232 | 7.81E-61 | 9.21E-60 | TBX1-related factors | 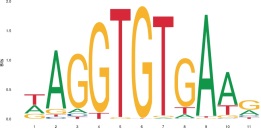 |
| 44 | MA0659.1_MAFG | 1.383708913 | 1.98E-59 | 2.28E-58 | Maf-related factors | 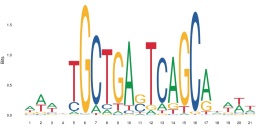 |
| 45 | MA0787.1_POU3F2 | 1.375917378 | 3.17E-57 | 3.58E-56 | POU domain factors | 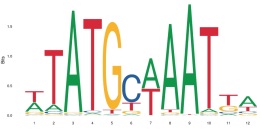 |
| 46 | MA0786.1_POU3F1 | 1.355048139 | 1.63E-51 | 1.79E-50 | POU domain factors | 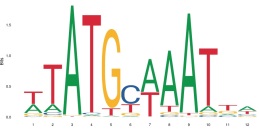 |
| 47 | MA0806.1_TBX4 | 1.346198148 | 3.51E-49 | 3.71E-48 | TBX2-related factors | 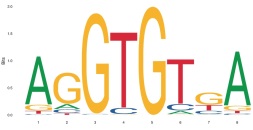 |
| 48 | MA0807.1_TBX5 | 1.346198148 | 3.51E-49 | 3.71E-48 | TBX2-related factors | 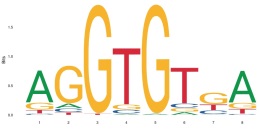 |
| 49 | MA0627.1_Pou2f3 | 1.339318795 | 2.11E-47 | 2.18E-46 | POU domain factors | 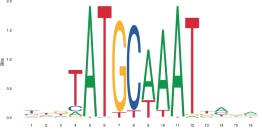 |
| 50 | MA0762.1_ETV2 | 1.336995215 | 8.28E-47 | 8.39E-46 | Ets-related factors | 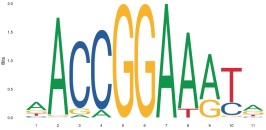 |
